# Nanoparticle Internalization Promotes the Survival of Primary Macrophages

**DOI:** 10.1101/2021.04.22.440822

**Authors:** Bader M. Jarai, Catherine A. Fromen

## Abstract

Macrophages, a class of tissue resident innate immune cells, are responsible for sequestering foreign objects through the process of phagocytosis, making them a promising target for immune-modulation via particulate engineering. Here, we report that nanoparticle (NP) dosing and cellular internalization via phagocytosis significantly enhances survival of *ex vivo* cultures of primary bone marrow-derived, alveolar, and peritoneal macrophages over particle-free controls. The enhanced survival is attributed to suppression of caspase-dependent apoptosis and is linked to phagocytosis and lysosomal signaling, which was also found to occur *in vivo*. Uniquely, poly(ethylene glycol)-based NP treatment does not alter macrophage polarization or lead to inflammatory effects. The enhanced survival phenomenon is also applicable to NPs of alternative chemistries, indicating the potential universality of this phenomenon with relevant drug delivery particles. These findings provide a framework for extending the lifespan of primary macrophages *ex vivo* for drug screening, vaccine studies, and cell therapies and has implications for any *in vivo* particulate immune-engineering applications.

## INTRODUCTION

Macrophages are leukocytes responsible for phagocytosing foreign objects, bacteria, and apoptotic cells, maintaining homeostasis, and bridging innate and adaptive immunities[1]. Hence, macrophages represent significant therapeutic targets and control of their phenotype, activity, and persistence could promote treatment for a wide range of conditions. Macrophages are highly responsive to their microenvironment and are equipped with phagocytic capabilities that enable them to internalize pathogens and larger particulates. Phagocytosis begins with ligand binding to cell surface receptors, triggering cytoskeletal rearrangement and actin polymerization to surround the target object. The target is then internalized within a phagosome, initiating a degradation process that proceeds through acidification of the compartment and lysosomal fusion[2]. Thus, the cell receives a plethora of information during phagocytosis that is critical to regulating cell function; however, decoupling the highly interconnected signaling pathways to precisely tune cell response remains an ongoing challenge.

Nano- and microparticles are natural avenues to stimulate macrophage function and elicit therapeutic response by leveraging these inherent phagocytic capabilities and delivering precision cues initiated during phagocytosis[3, 4]. Macrophages are highly sensitive to the cell-material interface, where everything from surface charge to particle shape can influence subsequent downstream signaling[5, 6]. Unsurprisingly, the role of material selection is critically important to tuning and directing these responses. Some materials used to fabricate nano- and microparticles have been shown to cause strong inflammatory or toxic effects on macrophages[7], while recent advances in biomaterials have elucidated classes of modular materials with strong biocompatibility[8–11]. Early generations of particulate therapeutics sought engineering solutions that avoided internalization by phagocytic immune cells to ensure successful cargo delivery[12, 13]; however, an emerging alternative is to promote controlled interactions with innate immune cells to regulate immune response[14, 15]. Indeed, particulates with no added stimulatory or tolerizing moieties show anti-inflammatory effects for preclinical treatment for West Nile virus, inflammatory bowel disease[16], sepsis[17], and acute lung injury[13, 18] that is presumably driven by particle phagocytosis and subsequent cell “distraction” of inflammatory phagocytes of monocytes and neutrophils, respectively.

Given the highly interconnected role between phagocytosis and macrophage phenotype, we sought to study downstream effects of primary macrophage internalization with “immunologically inert” poly(ethylene glycol) (PEG) diacrylate (PEGDA)-based nanoparticles (NPs), to uniquely investigate NP internalization in the absence of any known stimulatory cue, such as toll-like receptor (TLR) agonists or autophagy signals. Here, we report a surprising link between cell viability and particle phagocytosis in the absence of cell activation, allowing us to dramatically extend the viability of *ex vivo* macrophage cultures through particle treatment. We show that a single dose of NPs significantly delays primary macrophage cell death through the downregulation of caspase-dependent apoptotic pathways and activation of MAPK cascades, with no major changes to macrophage activation phenotype following NP uptake. Similar mechanisms are observed following *in vivo* administration, with the enhanced survival phenomenon occurring in *ex vivo* cultures of terminally differentiated macrophages isolated from tissue, as well as particles of alternative chemistries, indicating the potential universality of this phenomenon to any subset of macrophages with relevant drug delivery particles. Overall, this work uniquely demonstrates the ability of NPs to suppress apoptotic signaling and prolong viability in primary macrophages through phagocytosis signaling. This work could eliminate a major obstacle for macrophage-based cell immunotherapies and drug screenings that are typically hindered by poor macrophage survival, as well as provide mechanistic insight into the implications of NP phagocytosis on cell longevity for a wide range of therapeutic applications.

## RESULTS

### PEGDA NP phagocytosis stimulates primary macrophage survival *ex vivo*

As shown in **Figure 1A**, we observed that bone marrow-derived macrophages (BMMs) can persist in culture for up to 115 days following treatment with a single 20 μg/ml dose inert, unfunctionalized PEGDA NPs. This contrasts with untreated BMMs, which do not survive beyond a few weeks in culture. PEGDA NPs, approximately 300 nm in hydrodynamic size, were characterized using dynamic light scattering (DLS), scanning electron microscopy (SEM), and assayed for endotoxin levels that were determined to be non-significant (**Figure S1, S2**). To determine whether BMM survival is dependent on NP concentration, BMMs were incubated with a single dose of PEGDA NPs at concentrations ranging between zero and 100 μg/ml and survival was determined through metabolic activity as a measure of cell viability. From cell viability results, BMM survival was determined to be a strong function of NP concentration. One week following NP treatment, 20 μg/ml PEGDA NPs showed a 1.5x higher cell viability relative to untreated BMMs (p-value<0.05) while 50 and 100 μg/ml NP concentrations caused 3x and 3.5x higher cell viability compared to untreated BMMs (p-value<0.0001), respectively (**Figure 1B**). At a two-week timepoint, BMMs treated with 100 μg/ml and 50 μg/ml of NPs showed ~2.8x and 2.2x higher cell viability relative to untreated BMMs, respectively (p-value<0.0001), while the cell viability of BMMs treated with NPs at concentrations less than 50 μg/ml was statistically indistinguishable from cell viability of untreated BMMs. BMM count data were in agreement with cell viability as measured by metabolic activity and reflected strong NP concentration-dependent maintenance of BMM counts (**Figure 1C**). Furthermore, NPs treated to BMMs in the absence of fetal bovine serum showed no statistical differences in the resultant enhanced viability compared to BMMs in the presence of serum (**Figure S3**).

**Figure 1:**
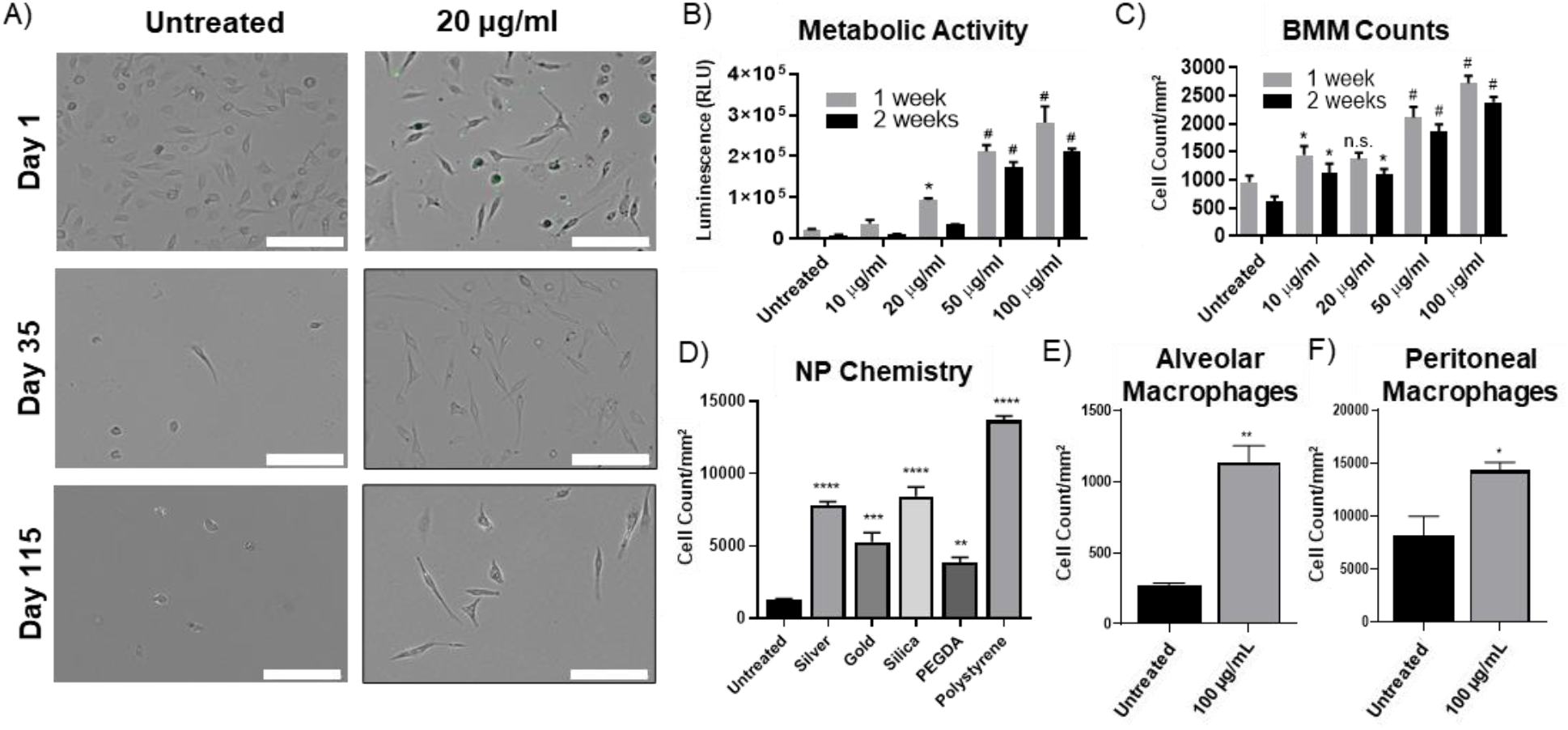
Phagocytosis of inert PEGDA NPs promotes longevity of primary macrophages *ex vivo*. A) Microscopy images at 20x magnification of BMMs treated with NPs (green fluorescence) showing enhanced survival compared to untreated cells for over 115 days. Scale bar 100 μm. B) Metabolic Activity of BMMs one and two weeks after treatment with different concentrations of PEGDA NPs. CellTiter-Glo® 2.0 Cell Viability Assay was used to determine cell viability measured through luminescence. *p<0.05, **p<0.01, #p<0.0001 (compared to untreated cells) using Dunnett’s multiple comparisons test as part of a two-way ANOVA. C) Cell counts of BMMs one and two weeks after treatment with different concentrations of PEGDA NPs. n.s. is not significant, *p<0.05, #p<0.0001 (compared to untreated cells) using Dunnett’s multiple comparisons test as part of a two-way ANOVA (*N*=6). D) Cell counts of BMMs treated with 100 μg/ml NPs of varying composition one week following NP treatment. *p<0.05, **p<0.01, ***p<0.001, ****p<0.0001 using Dunnett’s multiple comparisons test as part of a one-way ANOVA. E) Cell counts of *ex vivo* alveolar macrophages three weeks after treatment with 100 μg/ml PEGDA NPs. **p<0.01 using a student’s T-test. F) Cell counts of *ex vivo* peritoneal macrophages three weeks after treatment with 100 μg/ml PEGDA NPs. *p<0.05 using a student’s T-test. In all graphs, unless stated otherwise, bars represent the mean and error bars represent SEM (*N*=3); Data are representative of two independent experiments.

To investigate whether the PEGDA NP-driven enhanced survival is applicable to NPs of varying compositions, BMMs were dosed with 100 μg/ml silver, gold, silica, and polystyrene NPs and cell counts were used to evaluate survival after one week. All the tested NPs resulted in statistically significantly higher cell counts than untreated BMMs (**Figure 1D)**. BMMs treated with polystyrene NPs had the highest counts and more than 10x the counts of untreated BMMs with p-value<0.0001 using Dunnett’s multiple comparisons test as part of a one-way ANOVA. Silica and silver NPs resulted in approximately 6x higher cell counts compared to untreated BMMs with p-value<0.0001. BMMs treated with gold NPs had approximately 4x higher numbers than untreated cells with p-value<0.001 while PEGDA NPs showed the smallest improvement yet statistically significant with p-value<0.01 and around 3x higher counts of PEGDA NP-treated BMMs compared to untreated cells. All NP stocks were evaluated for the presence of endotoxin (**Figure S2**), with only silver and polystyrene detectable over the baseline. Overall, these results show that enhanced survival is possible with many NP formulations and are not restricted to PEGDA or polymeric NPs.

NP-viability enhancement to *ex vivo* survival was not restricted to BMMs. Murine alveolar macrophages treated with a single dose of 100 μg/ml NPs on day 1 show significantly higher counts than untreated cells (p-value<0.01) three weeks following isolation from murine lungs (**Figure 1E**). Similarly, peritoneal macrophages treated with 100 μg/ml of NPs display *ex vivo* longevity three weeks following isolation and a single dosage on day 1 (p-value<0.05) (**Figure 1F**). Results from NP-driven longevity of alveolar and peritoneal macrophages combined with those of BMMs highlight the versatility of utilizing NPs for enhancing the *ex vivo* survival of any macrophage type.

### NP-dependent survival is not dependent on macrophage polarization

Flow cytometric analysis of M1 and M2 markers was executed on BMMs of different initial polarization states 24 hours following NP treatment (Representative gating of flow cytometric data in **Figure S4**). No significant changes were observed in expression of either representative M1 markers (CD38, CD86) or M2 markers (CD206, EGR2) in NP-treated BMMs (**Figures S5A-D)** according to a student’s T-test (p-value>0.05) indicating no considerable M1 or M2 polarization of BMMs because of NP phagocytosis. Further, pre-skewed M1 and M2 BMMs treated with NPs did not display any significant change in CD38, CD86, CD206, and EGR2 expression, indicating that PEGDA NP phagocytosis promotes the survival of macrophages *ex vivo* without affecting their polarization state. Increased expression of MHCII was observed in M0, M1, and M2 BMMs (**Figure 2A**), indicating the potential enhancing of antigen presentation in NP-treated macrophages of all the tested phenotypes.

**Figure 2:**
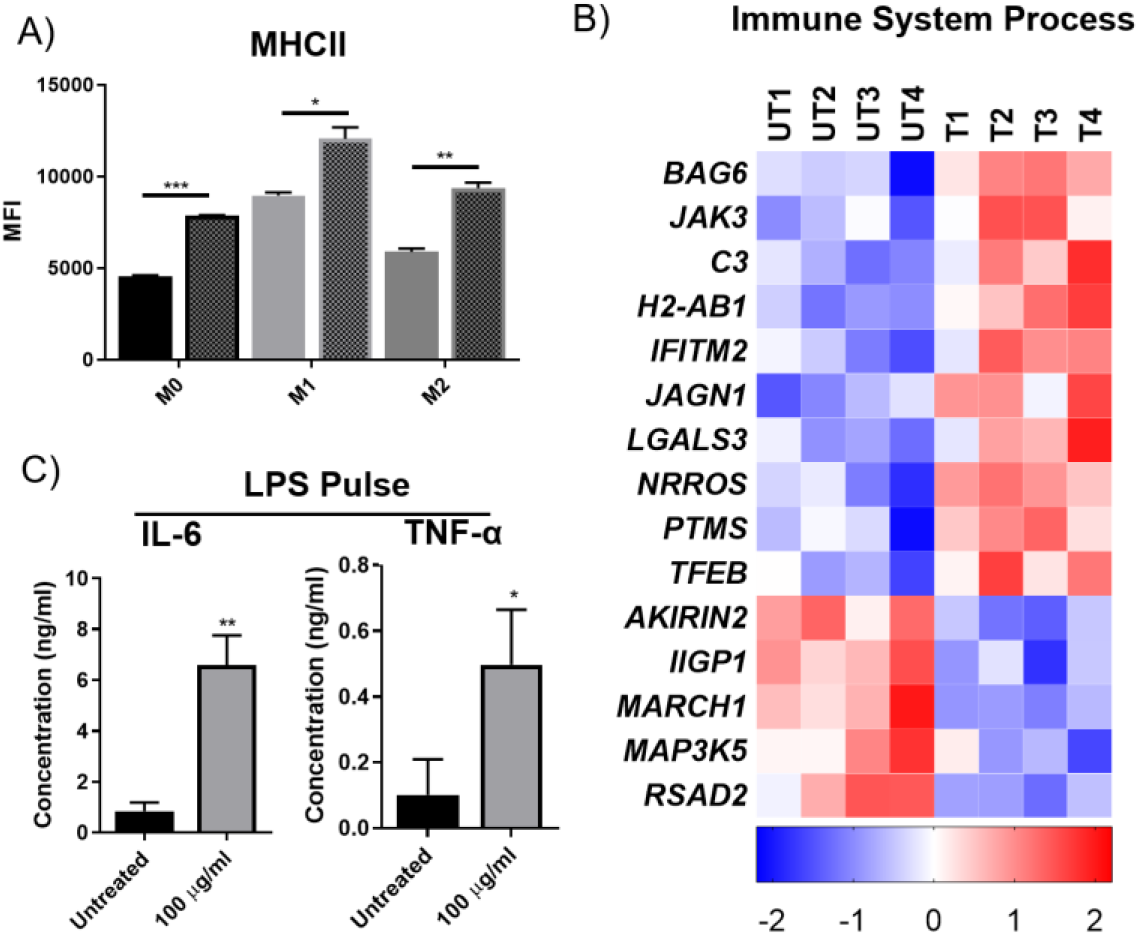
Phagocytosis of PEGDA NPs promotes macrophage immune functionality. A) MHCII marker expression of M0, M1, and M2 BMMs 24 hours following treatment with 100 μg/ml NPs (patterned bars). Data are representative of two independent experiments. B) Row Z-scores of gene counts from RNAseq analysis in Immune System Process GO term of NP-treated BMMs (T) compared to untreated BMMs (UT) showing four biological replicates per group. C) IL-6 and TNF-α concentrations of BMM supernatants four week following NP treatment after LPS challenge for 24 hours. In all plots, bars represent the mean and error bars represent SEM (*N*=3). *p<0.05, **p<0.01, ***p<0.001 using student’s T-test.

RNA sequencing (RNAseq) analysis was performed to identify relevant differentially expressed genes (DEGs) and perform functional classifications and clustering to determine significant cell processes involved in cell-NP interactions that ultimately lead to enhanced survival. Gene expression in BMMs treated with 100 μg/ml NPs for 24 hours was compared to that of untreated BMMs under the same culture conditions. Of the DEGs analyzed, 331 were determined to be statistically significant with q-value<0.05, but fold change values were relatively low, ranging between −2 and 2. The database for annotation, visualization and integrated discovery (DAVID), was used to identify enriched gene ontology (GO) terms in the 331 significant DEGs. Full DAVID enrichment analysis of GO terms is represented in **Table S1** (Selected significant GO terms in **Figure S6).** DEGs in the Immune System Process GO term were used to guide our subsequent analysis. RNAseq analysis confirms MHCII upregulation (*H2-AB1* gene) and points to a small panel of 15 genes involved in immunity and regulating immune cell function (**Figure 2B**).

However, potent changes to characteristic inflammatory genes were notably absent from this panel, which confirms conclusions from flow cytometric analyses of polarization markers showing little effect of NP treatment on macrophage activation phenotype. Further multiplex analysis of M0 macrophage supernatants for a broad array of cytokines and chemokines only identified detectable levels of IP-10, KC, MCP-1, MIP-2, VEGF, TNF-α in supernatants of untreated or NP-treated BMMs (**Figure S5E**). However, no significant change in any cytokine levels in BMMs treated with 100 μg/ml PEGDA NPs was observed compared to their untreated counterparts as determined by student’s T-tests (p>0.05). These results support conclusions from flow cytometric analyses indicating no notable impact of PEGDA NP phagocytosis on macrophage activation phenotype and that the NP-induced macrophage longevity does not require macrophage activation and/or classical polarization pathways.

While macrophage polarization was not required to observe the enhanced viability, long-living NP-treated cells were observed to respond appropriately to immune stimuli. As shown **Figure 2C** we observed that long living macrophages treated with 100 μg/ml PEGDA NPs were able to provide response to LPS challenge four weeks following NP treatment. Treated cells produced expected cytokines of IL-6 and TNF-α (**Figure 2C)** in culture supernatants that are statistically significant compared to untreated cells (as determined by student’s T-tests p-values<0.01 and 0.05, respectively).

### PEGDA NP treatment suppresses apoptotic pathways and promotes pro-survival signaling in *ex vivo* macrophages

In accordance with literature, untreated BMMs were found to undergo apoptosis[19], or programed cell death, that was delayed following NP treatment. Quadrant analysis after 24 hours in culture revealed rapid cell death through apoptosis of untreated cells (**Figure 3A, S7**). Treatment with 100 μg/ml NPs showed retardation of all major quantified modes of cell death, which was corroborated by suppression of active pro-apoptotic caspase 3/7 expression, an indicator of early apoptosis, in BMMs treated with NPs in a concentration-dependent manner (**Figure 3B**). Three days following treatment with 100 μg/ml NPs resulted in a significantly lower active caspase 3/7 expression (p-value<0.01), while the margin of active caspase 3/7 reduction is diminished with lower NP concentrations. TUNEL analysis revealed DNA damage only in untreated cells, which occurs during the late stages of apoptosis. At 24 hours NP-treated BMMs showed considerably lower fluorescent TUNEL signal than untreated BMMs (**Figure 3C**), indicating lower late apoptosis characterized by DNA damage in BMMs treated with NPs, which is also supported by analysis at 72 hours (**Figure S8**).

**Figure 3:**
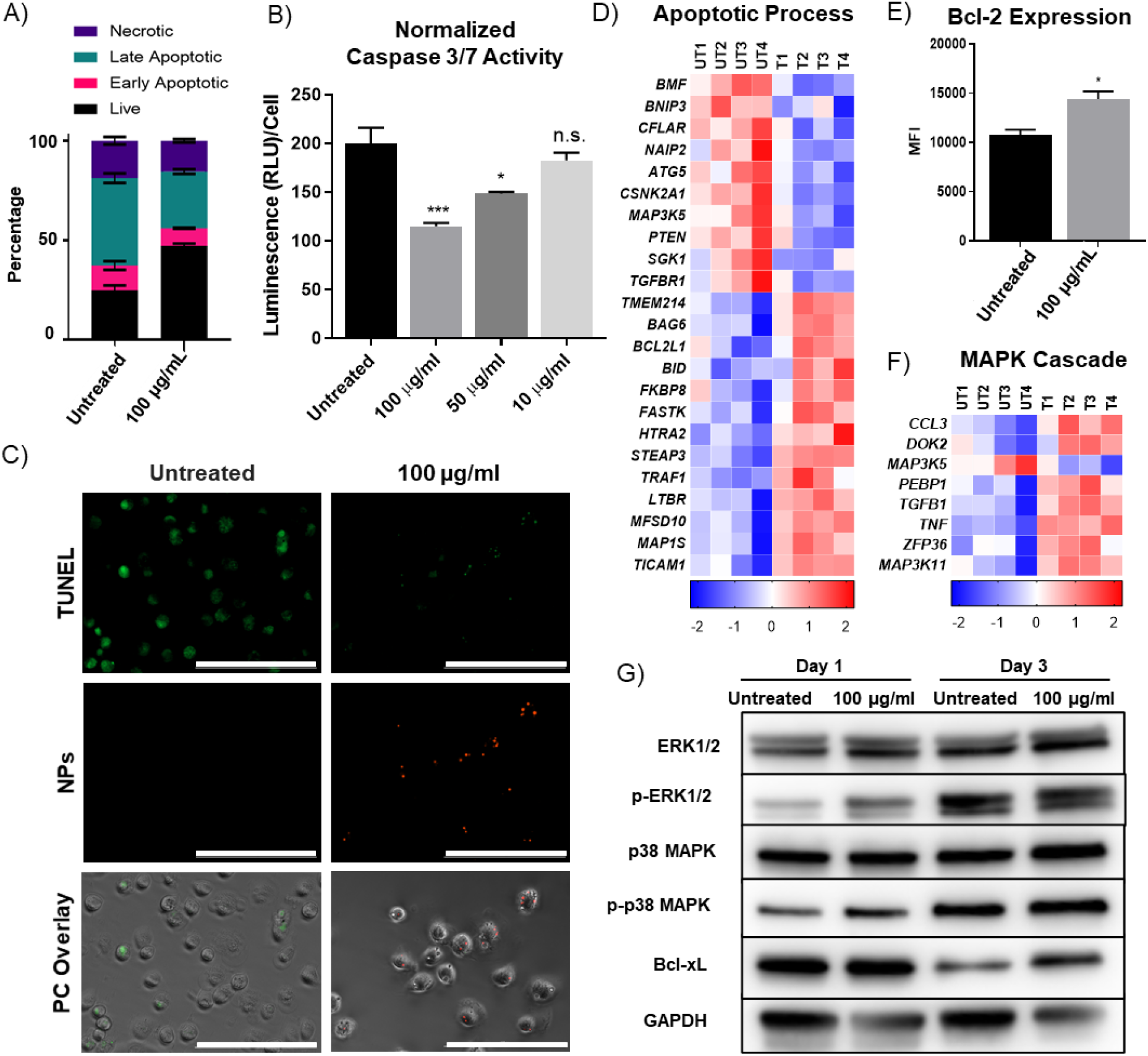
Treatment with PEGDA NPs suppresses apoptosis in primary macrophages. A) Representative flow cytometric quadrant analysis of modes of cell death in BMMs 72 hours following NP treatment based on Annexin V and Zombie Yellow dual staining. B) Normalized caspase 3/7 activity measured using Caspase-Glo® 3/7 Assay System 72 hours following NP treatment of varying concentrations. n.s. is not significant, *p<0.05, ***p<0.001 (compared to untreated cells) using Dunnett’s multiple comparisons test as part of a one-way ANOVA. Bars represent the mean and error bars represent SEM (*N*=3). C) TUNEL apoptosis imaging analysis of untreated or NP-treated BMMs 24 hours following NP treatment at 40x magnification. Scale bar 100 μm. Phase Contrast (PC). D) Row Z-scores of gene counts from RNAseq analysis in Apoptotic Process GO term of NP-treated BMMs (T) compared to untreated BMMs (UT) showing four biological replicates per group. E) Anti-apoptotic Bcl-2 protein expression 72 hours following NP treatment, measured by median fluorescent intensity (MFI) using Flow Cytometry. *p<0.05 using a student’s T-test. Bars represent the mean and error bars represent SEM (*N*=3). F) Row Z-scores of gene counts from RNAseq analysis in MAPK Cascade GO term of NP-treated BMMs (T) compared to untreated BMMs (UT) showing four biological replicates per group. G) Western blots of representative antiapoptotic proteins and members of MAPK cascades 1 day and 3 days following NP treatment. Data in A, B, E, and G are representative of two independent experiments.

RNAseq results were further evaluated to provide insight into the factors contributing to apoptosis suppression. Analysis of RNAseq revealed upregulation of several anti-apoptotic genes in NP-treated BMMs including *BCL2-L1* (**Figure 3D)** indicating a cascade of anti-apoptotic signaling is involved in promoting survival of *ex vivo* cultures of macrophages following NP treatment. In addition, several pro-apoptotic genes were downregulated in NP-treated BMMs, most notably Bcl-2-modifying factor (*BMF)* gene, which provides additional evidence to support the ability of NPs to suppress apoptotic signaling in macrophages *ex vivo*. Furthermore, flow cytometric analysis on BMMs treated with 100 μg/ml NPs for 3 days revealed significant upregulation of expression of B cell lymphoma-2 (Bcl-2) anti-apoptotic protein relative to untreated BMMs (p-value<0.05), which indicates the potential involvement of Bcl-2 in regulating macrophage survival and longevity caused by NP phagocytosis (**Figure 3E**). To further understand the cellular mechanism by which NPs delay rapid apoptosis in *ex vivo* cultures of macrophages, we assessed the expression of B cell lymphoma-extra large (Bcl-xL), which is encoded by the *BCL2-L1* gene and is known to interact with and inhibit executioner caspases 3 and 7 to promote cell survival (**Figure 4E**)[20]. Bcl-xL expression was higher in lysates of BMMs treated with 100 μg/ml NPs compared to untreated BMMs, especially at the 3 day timepoint.

**Figure 4:**
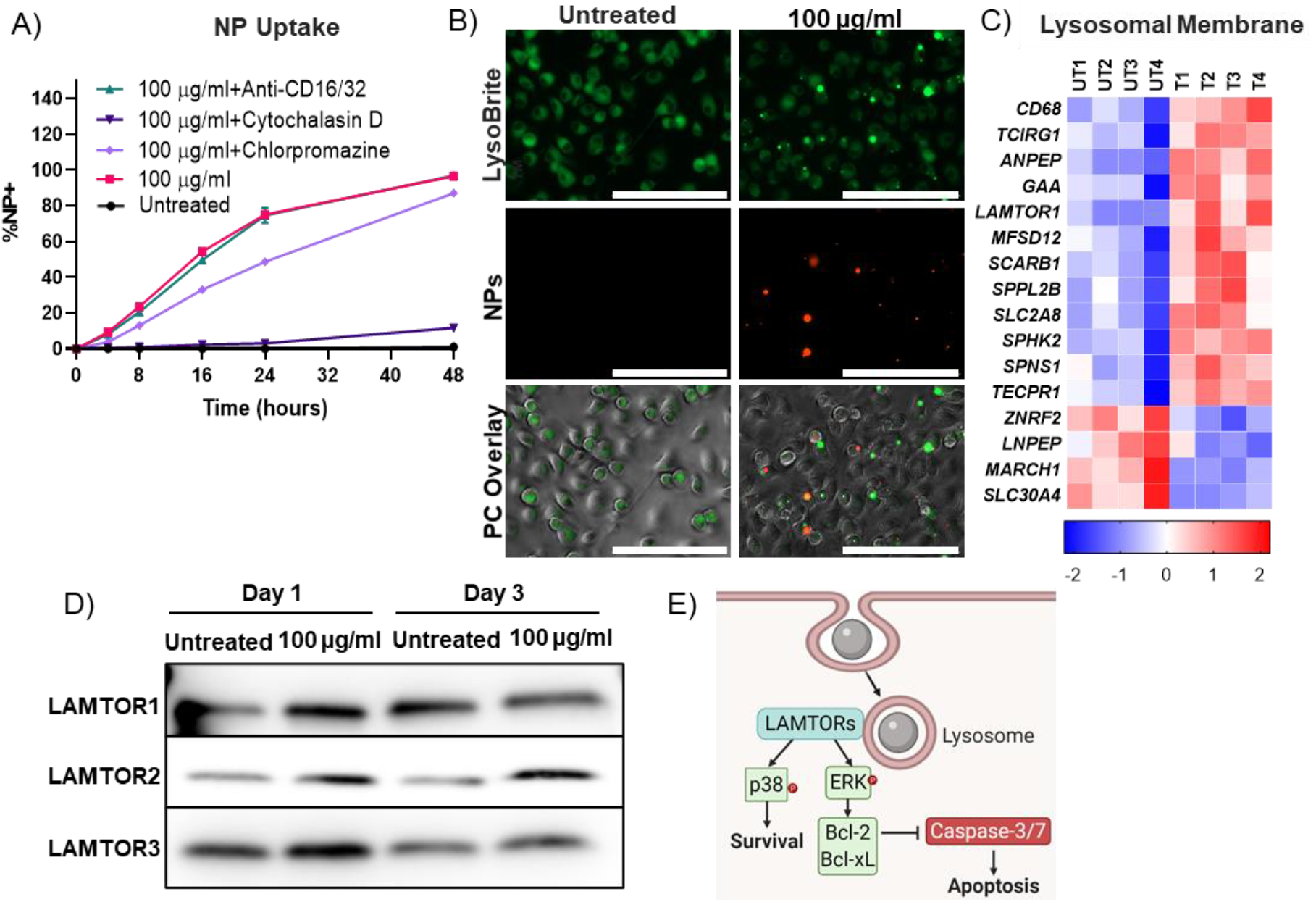
NP phagocytosis and lysosomal involvement in BMM survival. A) Kinetic NP uptake analysis and effect of internalization inhibitors on cell uptake. Error bars represent SEM (*N*=3). B) Row Z-scores of gene counts from RNAseq analysis in Lysosomal Membrane GO term of NP-treated BMMs compared to untreated BMMs showing four biological replicates per group. C) Lysosomal tracking and imaging at 40x magnification with LysoBrite™ Green 24 hours following NP treatment. Scale bar 100 μm. Phase Contrast (PC). D) Western blotting of LAMTOR1, 2, and 3 lysosomal proteins 1 day and 3 days following NP treatment. E) Schematic of hypothesized effect of NP phagocytosis on macrophage survival.

RNAseq analysis and enriched GO terms revealed the involvement of MAPK cascade genes, which are involved in cell function and survival (**Figure 3F**). Notably, we observed the upregulation of *MAP3K11* gene, which has been shown to interact with ERK and other kinases that regulate cell survival[21], and the downregulation of *MAP3K5* gene, which is known for its pro-apoptotic role especially in some disease environments[22, 23]. Guided by RNA sequencing analysis showing the involvement of MAPK cascade in NP-treated cells and the role of MAPK proteins in regulating cell survival and suppressing apoptosis[24], we elected to determine whether proteins in the MAPK cascade are involved in NP-induced macrophage longevity. Analysis of Western blots showed activation of p38 MAPK through increased phosphorylation in NP-treated BMMs compared to their untreated counterparts (**Figure 3G**). In addition, strong phosphorylation of ERK 1 and 2 (MAPK p42 and p44) was revealed by Western blotting in NP-treated BMMs (**Figure 3G**). These results confirm the increased involvement of MAPK cascade kinases and their activation following NP treatment.

### Phagocytosis and intracellular trafficking to lysosomal membranes is involved in *ex vivo* NP-treated macrophages with links to MAPK pathway activation and survival

NP entry into the cell can occur through several internalization routes, each contributing to different cellular signaling pathways[4]. Uptake of Cy5-labeled NPs in the presence or absence of inhibitors of internalization was assessed kinetically over a 48-hour period (**Figure 4A**). In the absence of inhibitors, approximately 75% of cells were positive for NPs within 24 hours and approximately 95% of cells were particle bearing by 48 hours. Pretreatment with anti-CD16/32 antibodies, which block Fc-mediated internalization did not result in considerably lower NP uptake with %NP+ cells closely matching those without the antibodies at 24 and 48 hour timepoints. Incubation with chlorpromazine, an internalization inhibitor especially for clathrin-mediated endocytosis, resulted in moderately lower uptake with 48.7% ± 2.7% and 87.1% ± 1.4% NP+ cells at 24 and 48 hours, respectively. A drastic drop in uptake resulted upon pretreatment with cytochalasin D, a potent inhibitor of actin-dependent internalization pathways especially phagocytosis, with 3.1% ± 0.2% and 11.7% ± 1.7% NP+ cells at 24 and 48 hours, respectively. These results indicate that actin-dependent internalization and especially phagocytosis was dominant in BMMs treated with PEGDA NPs.

Enrichment analysis of RNAseq results using DAVID revealed noteworthy involvement of 16 lysosomal membrane genes differing between untreated and treated cells (p-value<0.0001) (**Figure 4B**). *CD68* gene was notably upregulated in NP-treated BMMs. *CD68* encodes a transmembrane glycoprotein, which is a member of the lysosomal/endosomal-associated membrane glycoprotein (LAMP) and the scavenger receptor families and is involved in regulating phagocytosis and clearing debris[25]. In addition, *SCARB1* gene was upregulated, suggesting that NPs were phagocytosed by BMMs potentially through scavenger receptors and processed by involving lysosomal compartments. Furthermore, the upregulation of late endosomal/lysosomal adaptor, MAPK and mTOR activator 1 gene (*LAMTOR1*) was noteworthy, suggesting a link between phagocytosis and lysosomal membranes and activation of downstream cell signaling influencing survival. Lysosomal tracking through imaging with LysoBrite™ Green at 24 and 72 hours (**Figures 4C, Figure S9**) revealed high intensity regions of lysosomal activity in NP-treated BMMs compared to untreated controls, which may correspond to NP trafficking in late lysosomal compartments. Western blotting for LAMTOR proteins revealed a generally higher expression in NP-treated cells compared to the untreated control (**Figure 4D**). LAMTOR1 and LAMTOR3 protein expression was higher in lysates of BMMs treated with 100 μg/ml NPs compared to untreated BMMs especially at the day 1 timepoint. LAMTOR2 expression was consistently higher in NP-treated cells than in untreated cells both at day 1 and day 3 of analysis. Overall, these results strongly suggest that NP phagocytosis and subsequent lysosomal involvement are primary processes involved in cell-particle interactions that ultimately lead to enhancing survival.

### *In vivo* dosing of NPs and internalization stimulates the lysosome and upregulates pro-survival factors

100 μg of NPs were dosed to mice via orotracheal instillations and intraperitoneal injections to investigate the *in vivo* implications of NP phagocytosis. Alveolar and peritoneal lavages were performed at 24 hours following administration and CD11b+ macrophages were considered for analysis (Representative flow cytometric gating in **Figure S10**). NP uptake by macrophages was assessed (**Figure 5A**); approximately 83% and 80% of peritoneal and alveolar macrophages, respectively, were determined to be NP+ via flow cytometry. Lysosomal tracking in alveolar and peritoneal macrophages was performed through imaging with LysoBrite™ Green (**Figure S11**) and revealed high intensity regions of lysosomal activity in alveolar and peritoneal macrophages from NP-dosed mice compared to their untreated counterparts, which may be associated with NP trafficking to late lysosomal compartments and subsequent stimulation of lysosomal signaling.

**Figure 5:**
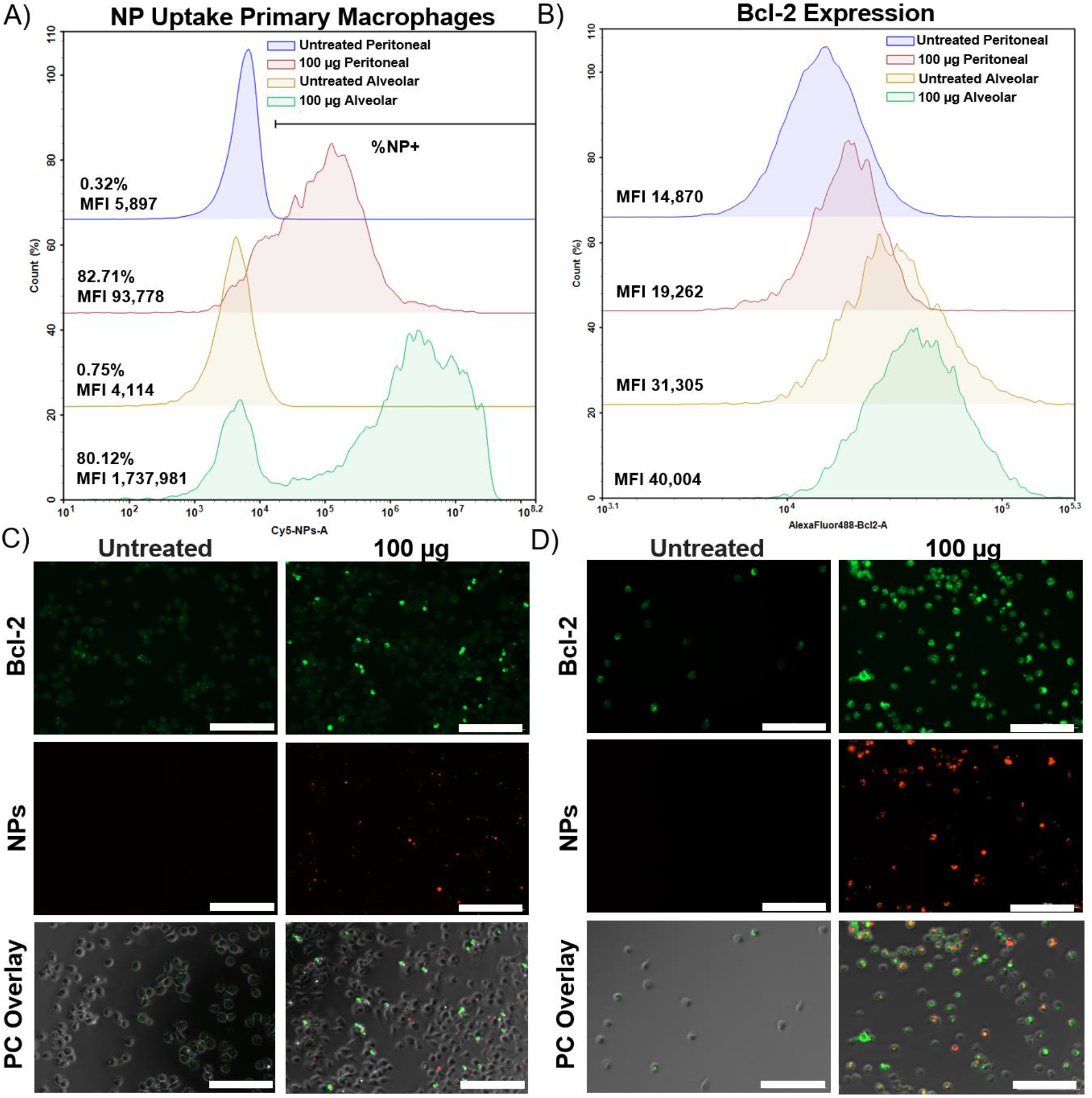
NP phagocytosis *in vivo* upregulates anti-apoptotic Bcl-2 protein in alveolar and peritoneal macrophages. A) Cy5-labeled NP internalization in peritoneal macrophages and alveolar macrophages. %NP+ cells and MFI (median fluorescence intensity) are indicated. B) Flow cytometric analysis of Bcl-2 expression in *in vivo* NP-treated peritoneal macrophages and alveolar macrophages. Data are representative of two independent experiments. Fluorescent imaging at 20x magnification after immunostaining with anti-Bcl-2 antibodies 24 hours following dosing of 100 μg of NPs to mice: C) Peritoneal macrophages D) Alveolar macrophages. Scale bar 100 μm. Phase Contrast (PC).

To investigate whether NP internalization by macrophages *in vivo* has pro-survival implications, anti-apoptotic Bcl-2 expression was evaluated in alveolar and peritoneal macrophages from NP-dosed mice. Flow cytometric analysis of alveolar and peritoneal macrophages from NP-dosed mice (**Figure 5B**) showed a markedly higher Bcl-2 expression in both alveolar and peritoneal macrophages from NP-dosed mice compared to their untreated counterparts. Median fluorescence intensities (MFI) of AlexaFluor488-conjugated anti-Bcl-2 antibodies were 14,870 and 19,262 for peritoneal macrophages from untreated and NP-dosed mice, respectively and 31,305 and 40,004 for alveolar macrophages from untreated and NP-dosed mice, respectively, indicating upregulation of anti-apoptotic Bcl-2 upon *in vivo* NP dosing. Immunostaining reveals notably higher expression of Bcl-2 in peritoneal and alveolar macrophages from NP-dosed mice relative to their untreated counterparts, characterized by higher intensity of green fluorescence (**Figure 5C,D**) confirming the results from flow cytometric analyses. Interestingly, TUNEL analysis revealed no notable differences between peritoneal macrophages from untreated and NP-dosed mice (**Figure S12**), indicating low overall apoptotic activity in *in vivo* macrophages despite NP-driven Bcl-2 upregulation.

## DISCUSSION

In this study, we demonstrated that dosing primary macrophages with inert and unfunctionalized NPs results in enhanced *ex vivo* survival for extended periods of time. Survival of macrophages was a strong function of NP concentration and did not rely on traditional activation and polarization pathways, which indicates a unique pro-survival effect in the absence of hallmark TLR stimulation. Macrophages dosed with NPs avoided apoptosis through mechanisms linked with phagocytosis and downstream NP processing in lysosomal compartments. Enhanced macrophage longevity was observed following treatment of a wide range of particle chemistries in BMMs, as well as primary alveolar and peritoneal macrophages, suggesting a universal effect of particle internalization by phagocytes driving cell lifespan. Furthermore, *in vivo* dosing of NPs to mice resulted in upregulation of pro-survival proteins and stimulation of lysosomal activity, in agreement with *ex vivo* results. This work has important implications for understanding and regulating cell lifespan through internalization of any foreign body.

BMMs are an established *in vitro* primary macrophage model that are differentiated *ex vivo* and are widely used to study macrophage function. BMMs require relatively high amounts of granulocyte-macrophage colony-stimulating factor (GM-CSF) for differentiation and proliferation, yielding high numbers of primary-like macrophage cells. However, BMMs rapidly undergo apoptosis in the absence of GM-CSF stimulating factors[19]. Our results demonstrated that a single dosage of PEGDA NPs in the absence of other signals, including GM-CSF, can dramatically extend their lifespan. In addition, we investigated the effect of NP dosing on the survival of terminally differentiated primary alveolar and peritoneal macrophages *ex vivo*. Our results demonstrated that treatment with PEGDA NPs significantly improved the survival of all the tested macrophage types, indicating that NP-induced survival may extend to a wide range of macrophage populations and phagocytic cells. The identified pathways linking phagocytosis, lysosomal signaling, and cell lifespan are ubiquitous to many phagocytes (**Figure 4E**); thus, similar extended viability would be expected for a wide range of innate immune cells following particulate internalization[26]. While our results have yet to explore additional phagocytes or their extended survival in the *in vivo* environment, we have demonstrated NP-driven upregulation of pro-survival Bcl-2 proteins and stimulation of lysosomal activity following *in vivo* NP dosing, indicating that particulate-induced cell survival may occur *in vivo*. Furthermore, poorly soluble particulate adjuvants (*e.g.,* aluminum hydroxide [alum], talc, oil-in-water emulsions, aggregated oxidized low density lipoproteins [ox-LDL]) have been shown to extend phagocyte survival (*i.e.,* macrophage, dendritic cell) at the site of injection in the absence of GM-CSF[27]. While this extended survival is often attributed to inflammatory signaling cascades or macrophage polarization, our results with non-stimulatory PEGDA NPs suggests survival can be independently promoted through phagocytosis and likely enhanced with inflammatory synergies. Further explorations in other phagocytes and macrophage populations both *ex vivo* and *in vivo* are warranted to continue to decouple these effects.

Our study leveraged PEGDA hydrogel NPs, commonly used in the drug delivery field to extend circulation times, encapsulate hydrophilic cargos, and evaluate physiochemical NP properties[28, 29]. Importantly, PEGDA nanoparticles have been shown to be immunologically inert[30], in contrast to commonly used biodegradable hydrophobic polyesters, such as poly(lactic-co-glycolic acid) (PLGA), that yield acidic degradation products with immunomodulatory effects[31]. Indeed, BMMs treated with PEGDA-NPs showed no evidence of PRR activation (**Figure S5**), allowing us to confirm the effects of NP-induced longevity occur independently from stimulatory signaling pathways. However, the NP-enhanced survival of BMMs was not found to be unique to this formulation; BMMs treated with other unfunctionalized NPs of different compositions all displayed enhanced viability following a single administration. Polystyrene, silica, gold, and silver NPs resulted in significantly higher BMM counts after one week relative to untreated BMMs and were all higher than cell counts of PEGDA NP-treated BMMs. We hypothesize that variable NP uptake contributes to differences in the resulting survival; relative to PEGDA NPs, other NPs are composed of stiffer materials which will promote higher uptake in macrophages[28]. Furthermore, silver, gold, silica, and polystyrene NPs have been shown to cause activation and polarization in macrophages[9], which could also explain differences in survival across nanoparticle types and points to synergize arising from inflammatory and phagocytic cascades. Regardless, the BMM response to the range of NP groups highlight that NP-induced macrophage survival may be universally applicable to NPs of all compositions, emphasizing the broad importance of our findings for the drug delivery community.

Within 24 hours of GM-CSF depletion, a significant portion of *ex vivo* BMMs transitioned to apoptotic regimes, supported by executioner caspase activity and DNA damage assays (**Figure 3**). Suppression of executioner caspase 3/7 activity is likely due to interactions with anti-apoptotic Bcl-2 family proteins including Bcl-2 and Bcl-xL, which were found to be upregulated in NP-treated *ex vivo* BMMs (**Figure 3**) as well as *in vivo*-dosed alveolar and peritoneal macrophages (**Figure 5**). Anti-apoptotic Bcl-2 family proteins have been known to promote survival through the inhibition of caspase activation in apoptosis signaling[20, 32]. Following phagocytosis, which we confirm is the dominate route of NP internalization (**Figure 4A**), internalized particles are channeled through cellular compartments starting from the phagosome and through the matured phagolysosome, triggering a myriad of signaling pathways[4]. Phagocytosis initiates through a multitude of surface receptors that can generally be grouped into two classes: opsonin-dependent (*i.e.* Fc and complement receptors, integrins) and opsonin-independent (*i.e.* scavenger receptors, c-type lectins)[26]. However, other internalization pathways may be responsible for NP uptake and trafficking including, *e.g.* pinocytosis and endocytosis, which shuttle the internalized NP to the lysosomal compartment. Regardless of the route of entry, the lysosome is a hotspot for signaling and is associated with a wide range of pathways regulating survival, including some MAPK pathway proteins (*e.g.,* ERK1/2) that are intimately intertwined with and regulated by lysosomal activity[33]. Indeed, RNAseq enrichment analysis (**Figure S6**) highlighted the role of MAPK, lysosomal proteins, and protein kinase B in the anti-apoptotic events. ERK1/2 activation has been linked to enhanced survival through the upregulation of anti-apoptotic proteins that belong to the Bcl-2 family (**Figure 4E**)[34]. Our results indicated that LAMTOR1, LAMTOR2, and LAMTOR3 were upregulated in cell lysates of NP-treated BMMs, corresponding to phosphorylation of ERK1/2 and p38 MAPK, in parallel to gene level upregulation of *LAMTOR1* and several genes in the MAPK cascade, which were ultimately reflected in the resulting anti-apoptotic activity. Depletion of lysosomal proteins, especially LAMTOR1 and LAMTOR2, has been shown to cause rapid apoptosis in phagocytes[35, 36], indicating the role of late endosomal and lysosomal cellular compartments in regulating survival. Our reports of increased LAMTOR activity (**Figure 4D**) implicated this family in macrophage resistance to apoptosis following NP phagocytosis. Along with ERK1/2 phosphorylation, protein kinase B and p38 MAPK activation have been shown to be involved in promoting cell survival[37]. Phosphorylation of p38 MAPK revealed an additional route through which NP phagocytosis may influence macrophage survival. Activation of p38 MAPK and its associated pathways has been implicated in apoptosis resistance in monocytes and macrophages[38]. Altogether, this unravels unexplored interactions of phagocytosis and survival signaling that are likely stemming from the involvement of lysosomal signaling.

Macrophage survival in response to external stimuli has been commonly linked to traditional TLR signaling leading to potent Nuclear Factor (NF)-κB activation and the transcription of inflammatory genes[39], which results in the secretion of cytokines and ultimately leading to polarization into an inflammatory phenotype. Stimulation of NF-κB signaling has been previously utilized to prolong the survival of *ex vivo* macrophages and other leukocytes using cytokines[40], TLR agonists,[39] adjuvants[41], and some lipoproteins (ox-LDL)[42]. Similar extension of phagocyte survival has also been attributed to autophagy signaling, with internalization of apoptotic cells promoting phagocyte survival through Protein kinase B (Akt) activation and MAPK ERK1/2 inhibition[43]. Autophagy can also regulate phagocyte activation state through NF-κB signaling degradation and the mechanistic target of rapamycin (mTOR) pathway, which has been attributed to M1 and M2 polarization[44, 45]. However, owing to the immunologically inert nature of PEG-based materials[30], PEGDA NPs did not lead to macrophage activation or significant changes to inflammatory cytokine secretions, which enabled the decoupling of survival and activation phenomena with respect to understanding the impact of NP phagocytosis on primary macrophage fate. Interestingly, PEGDA NP phagocytosis promoted upregulation of MHCII despite insignificant changes to M1 or M2 activation markers. While MHCII expression traditionally accompanies other polarization markers along the M1-M2 paradigm, including CD86 or CD206 in response to polarizing stimuli[46], these were not observed in our results (**Figure S5**). Other NP systems have been shown to cause MHCII upregulation, but these reports are often coupled with markers of macrophage polarization[9]. Thus, treating with PEGDA NPs could enhance MHCII expression for vaccine studies, *in vitro* drug screening, or immune-skewing therapeutics without significant impact on phenotypical state. This may be especially critical for developing NP-based tolerizing therapies, where presentation of antigens in the absence of pro-inflammatory stimuli may enhance the tolerogenic effects of the NP therapy, especially for treatment of allergic responses or autoimmune diseases[47].

The discoveries made in this work outline the strong dependence of macrophage survival on NP phagocytosis in *ex vivo* cultures, which are plagued by rapid apoptosis. These findings are of special importance for improving macrophage utility in drug discovery and screening, as well as biomanufacturing opportunities. Furthermore, autologous macrophage therapies[48], which commonly rely on *ex vivo* stimulation prior to *in vivo* administration will greatly benefit from increased survival throughout the stimulation period of the explant, which may extend to *in vivo* implants. In addition to more certain *ex vivo* survival improvements, the presented findings also indicate potential enhancement of macrophage survival in *in vivo* environments as a result of NP-driven upregulation of anti-apoptotic proteins. Both peritoneal and alveolar macrophages exhibited lysosomal involvement following *in vivo* NP administration that led to increasing Bcl-2 expression (**Figure 5**), in line with *ex vivo* results from these same primary cell types. Interestingly, alveolar macrophages expressed higher basal levels of Bcl-2 when compared to peritoneal macrophages, which may be attributed to differential function, exposure conditions, and lifespans; alveolar macrophages reside at a mucosal interface and can span many months to years[49], while peritoneal macrophages are less exposed to external stimuli and can present both short- and long-half-lives depending on their precursor origin[50]. Regardless, both cell types exhibited increased anti-apoptotic Bcl-2 expression following *in vivo* NP treatment, suggesting NP internalization *in vivo* will influence downstream cell viability. Upregulation of anti-apoptotic factors *in vivo* presents unique therapeutic opportunities where pro-apoptotic potential is imposed, for example in the case for some bacterial infections[51], where NP treatment to macrophages may suppress apoptosis and aid in host defense. In the context of drug delivery applications, the fate of NPs following *in vivo* administration is overwhelmingly affected by phagocytes, especially macrophages[4], NP interactions with phagocyte survival must be studied in drug delivery systems especially where macrophages may play a crucial role in the pathology, as is the case with atherosclerosis[52], asthma[53], or idiopathic pulmonary fibrosis[54]. Long-term studies are needed to uncover what, if any, negative effect increased survival may have on other cell functionality. Thus, thorough investigations of NP interactions with macrophages and other phagocytes are critical to understanding of the role of phagocytes in healthy and disease conditions and, ultimately, NP therapeutic employment.

Altogether, enhancing survival of macrophages leveraging NP platforms opens the door to a wide range of opportunities in therapeutic interventions with implications in drug delivery, drug screening, cell therapies, immune engineering. Furthermore, these results aid to decouple the mechanisms involved in phagocytosis pathways that regulate cell viability to promote predictive understanding of phagocyte lifespan both *ex vivo* and *in vivo*.

## METHODS

### Primary Macrophage Isolation and Murine Studies

All studies involving animals were performed in accordance with National Institutes of Health guidelines for the care and use of laboratory animals and approved by the Institutional Animal Care and Use Committee (IACUC) at the University of Delaware. C57BL/6J (Jackson Laboratories) were housed in a pathogen-free facility at the University of Delaware. Female C57BL/6J six to ten weeks of age were used to extract primary cells. BMMs and alveolar and peritoneal macrophages were isolated from mice according to standard protocols as described in the following.

To generate bone marrow-derived macrophages (BMMs), standard protocols as previously reported were followed[55]. Briefly, bone marrow was extracted from femurs and tibias of mice following euthanasia and cells were plated in BMM differentiation media (DMEM/F-12 media (Corning) containing 20% fetal bovine serum, 30% L929 cell conditioned media, and 1% Penicillin-Streptomycin). On day three, cells were supplemented with an additional dose of BMM differentiation media and used on day seven for experiments following removal of L929 cell conditioned media and culture in DMEM/F-12 media containing 10% fetal bovine serum.

For *in vivo* studies, 100 μg NPs in PBS were administered to mice to the lungs via an orotracheal instillation or to the peritoneal cavity via an intraperitoneal injection according to standard methods. Primary alveolar macrophages were extracted from mice by performing a bronchoalveolar lavage (BAL) and extracting the BAL fluid (BALF). Briefly, BALF was spun down in a cold centrifuge at 1500 rpm for five minutes and the cell pellet was resuspended in red blood cell lysis buffer (Invitrogen) for 60 seconds and then resuspended in DMEM/F-12 media containing 10% fetal bovine serum, and 1% Penicillin-Streptomycin and seeded in well plates for experiments. Primary peritoneal macrophages were isolated from mice by performing a peritoneal lavage (PL) and extracting the PL fluid (PLF) as previously described[55]. The PLF was spun down in a cold centrifuge at 1500 rpm for 5 minutes and the cell pellet was resuspended in DMEM/F-12 media containing 10% fetal bovine serum, and 1% Penicillin-Streptomycin and plated in well plates overnight to allow for peritoneal macrophages to adhere. Following overnight incubation, nonadherent cells were removed and cells were scraped and re-plated for experiments.

### NP Synthesis

Poly(ethylene glycol) diacrylate (PEGDA)-based NPs were synthesized as previously described[56]. Briefly, PEG_700_DA (M_n_ = 700), 2-carboxyethyl acrylate (CEA), and photoinitiator diphenyl(2,4,6-trimethylbenzoyl) phosphine oxide were dissolved in methanol (50wt% solution) and emulsified with silicone oil AP1000 (All from Millipore Sigma) (**Figure S1**) and polymerized by irradiating with UV light (365 nm at ∼28 cm from the light source, ∼5–10 mW/cm^2^) for the 30 seconds. Micron-sized particles were eliminated by passing the suspension through a series of sterile five-micron and one-micron filters. The resulting NPs were washed twice in sterile water prior to a final suspension in DMEM/F-12 media containing 10% fetal bovine serum, and 1% Penicillin-Streptomycin for use in experiments. In experiments requiring particle tracking, FITC (Millipore Sigma) or Cy5 maleimide (AAT Bioquest) fluorescent labels were included in the monomer mix at compositions of 1% and 0.25%, respectively. In experiments with non-PEGDA NPs, unfunctionalized 100 nm silver, gold, and silica NPs (Nanocomposix) as well as 100 nm polystyrene NPs (Millipore Sigma) were obtained.

### NP Characterization

Thermogravimetric analysis (TGA) using TA Instruments TGA 550 was utilized to determine NP concentrations following the final water wash. 50 μl of NP suspensions were added to TGA sample pans in technical triplicates at a series of dilutions. The mass of the NPs was determined via a mass reading after a temperature ramp to 120 °C followed by a 30-minute isothermal step to ensure water evaporation. Dynamic light scattering (DLS) was performed using a Malvern Zetasizer Nano S to characterize the hydrodynamic diameter of NPs. NPs were prepared for DLS by diluting to 0.1 mg/ml in deionized water. Hydrodynamic diameters and polydispersity indices were measured for technical triplicates. Scanning electron microscopy (SEM) was performed using a JSM F7400 scanning electron microscope following gold/palladium sputter coating using a Denton Desk IV sputter coater. Endotoxin contamination was assessed for all NP groups using Pierce™ LAL Chromogenic Endotoxin Quantitation Kit (ThermoFisher Scientific) according to manufacturer’s guidelines.

### NP Internalization and Trafficking

BMMs were seeded in 24-well plates (2.5×10^5^ cells/well) and allowed to adhere for at least four hours prior to NP treatment. NPs were administered to cells at the indicated final concentrations for 24 hours. Kinetic NP uptake was determined by dosing BMMs with 100 μg/ml Cy5-labelled NPs. Cells were detached using Accutase® (Innovative Cell Technologies, Inc.) at 0, 4, 8, 16, 24, and 48 hours and analyzed for %Cy5+ cells using ACEA NovoCyte Flow Cytometer. To differentiate internalization pathways, BMMs were treated with 2 μg/ml anti-CD16/32, 1.5 μg/ml Cytochalasin D, or 5 μg/ml Chlorpromazine hydrochloride 30 minutes prior to NP dosing and analyzed at 0, 4, 8, 16, 24, and 48 hours following NP treatment for %Cy5+ cells using Flow Cytometry. For lysosomal imaging, cells were cultured in glass bottom, black walled 96-well plates (0.5-1.5×10^4^ cells/well) and Cell Navigator™ Lysosome Staining Kit (AAT Bioquest) was used according to manufacturer’s guidelines. Cells were imaged using BioTek Cytation 5 Multimode Imager.

### Cell Viability Assessment

BMMs were seeded in 96-well plates (1×10^5^ cells/well) and allowed to adhere for at least four hours prior to NP treatment. NPs were administered to cells at the indicated concentrations for 24 hours. Metabolic activity as a measure of cell viability was assessed using CellTiter-Glo® 2.0 Cell Viability Assay (Promega) according to manufacturer’s guidelines. BioTek Cytation 5 Multimode Imager was also used to monitor cell counts as a measure of viability. Zombie Yellow™ Fixable Viability Kit (Biolegend) was used according to manufacturer’s guidelines for flow cytometric assessment of cell viability. For apoptosis detection, Caspase-Glo® 3/7 Assay System (Promega) was used according to manufacturer’s guidelines. Phosphatidylserine membrane translocation was quantified using fluorescent staining with Annexin V-Pacific Blue (Biolegend). Bcl-2 anti-apoptotic protein expression was measured by intracellular staining of BMMs with PE-Cy7 anti-Bcl-2 antibody (Biolegend). AlexaFluor488 anti-Bcl-2 and BV711 anti-CD11b antibodies (Biolegend) were used for staining of *in vivo* NP-treated alveolar and peritoneal macrophages. For TUNEL assay, cells were cultured in glass bottom, black walled 96-well plates (0.5-1.5×10^4^ cells/well) and Cell Meter™ Live Cell TUNEL Apoptosis Assay Kit (AAT Bioquest) was used according to manufacturer’s guidelines. Gene and protein analysis of NP-treated BMMs is described in the following sections.

### Western Blotting

BMMs were seeded in 6-well plates (1×10^6^ cells/well) and allowed to adhere for at least four hours prior to NP treatment. NPs were administered to cells at the indicated final concentrations for 24 hours. BMMs were harvested and lysed with ice-cold RIPA lysis buffer (Alfa Aesar) supplemented with Halt Protease and Phosphatase Inhibitors (ThermoFisher Scientific). Cell lysates were spun down at 16,000 RCF for ten minutes in a precooled centrifuge and supernatants were collected. Protein content of supernatants was determined using a BCA assay (ThermoFisher Scientific). Following denaturation of protein samples in Laemmeli sample buffer (Bio-Rad), 15 μg of protein were loaded onto 4-20% Mini PROTEAN gels (Bio-Rad) and run at 50 V for five minutes followed by 150 V for 60 minutes in running buffer (25 mM Tris, 192 mM glycine, and 0.1% SDS, pH 8.3). Protein bands were transferred to PVDF membranes (Bio-Rad) and blocked for two hours with 5% bovine serum albumin (BSA) in Tris-buffered saline with 0.1% Tween20 (TBST) and incubated overnight in Bcl-xL, p44/42 MAPK (ERK1/2), phospho-p44/42 MAPK (ERK1/2) (Thr202/Tyr204), GAPDH, LAMTOR1, LAMTOR2, LAMTOR3, p38 MAPK, phospho-p38 MAPK(Thr180/Tyr182)anti-mouse primary antibodies diluted according to manufacturer’s guidelines in TBST containing 5% BSA. Membranes were then washed three times in TBST and incubated with horseradish peroxidase-conjugated anti-Rabbit IgG secondary antibody (All antibodies from Cell Signaling Technology) at 1:2000 dilution in TBST containing 5% BSA. Membranes were then washed three times in TBST prior to incubation with Amersham Chemiluminescent Detection Set (GE Healthcare) for visualization. Membranes were imaged using Azure 280 Imager. Each membrane was reserved for each primary antibody and all the reported blots are from the same batch of lysates; housekeeping protein (GAPDH) was blotted independently for the same batch of lysates.

### Macrophage Polarization Studies

BMMs in 6-well plates (1×10^6^ cells/well) were detached using Accutase® (Innovative Cell Technologies, Inc.) and washed twice with PBS supplemented with 2% FBS. Cells were then blocked with anti-CD16/32 (Fc block, Biolegend) for 10 minutes and then surface-stained for 30 minutes with the following antibodies: CD38-Pacific Blue, CD86-AlexaFluor700, and I-A/I-E-Brilliant Violet 785™ (All from Biolegend). Cells were then fixed with 4% paraformaldehyde in PBS (Alfa Aesar) for 15 minutes and then permeabilized using Intracellular Staining Permeabilization Wash Buffer (Biolegend) and stained for intracellular markers with the following antibodies: CD206-PE-Cy7 and EGR2-APC (All from Biolegend) and analyzed using ACEA NovoCyte Flow Cytometer. Median fluorescent intensity (MFI) was recorded via flow cytometry as a measure of marker expression. In some studies, 25 ng/ml lipopolysaccharides (LPS) from *Escherichia coli* O111:B4 (Millipore Sigma) or 25 ng/ml Interleukin-4 (IL-4) (Biolegend) were used to stimulate M1 or M2 polarization, respectively for 24 hours prior to use in experiments. For cytokine analyses, concentrations of supernatant at no dilution were measured in BMM supernatants by the Cytokine Core Laboratory at the University of Maryland (Baltimore) for Eotaxin, G-CSF, GM-CSF, IFN-γ, IL-1α, IL-1β, IL-2, IL-3, IL-4, IL-5, IL-6, IL-7, IL-9, IL-10, IL-12 p40, IL-12 p70, IL-13, IL-15, IL-17α, IP-10, KC, LIF, LIX, MCP-1, MCSF, MIG, MIP-1α, MIP-1β, MIP-2, RANTES, TNF-α, and VEGF-α using a 32-panel LUMINEX multianalyte system. Enzyme-Linked Immunosorbent Assays (ELISAs) kits (BD Biosciences) for Interleukin-6 (IL-6) and Tumor Necrosis Factor-α (TNF-α) were also performed on BMM supernatants according to manufacturer’s guidelines.

### RNA Sequencing and Bioinformatics

BMMs were seeded in 6-well plates (1×10^6^ cells/well) and allowed to adhere for at least four hours prior to NP treatment. NPs were administered to cells at the indicated final concentrations for 24 hours. BMMs were lysed *in situ* and RNA was extracted and purified using RNAeasy Plus Mini Kit (Qiagen) according to manufacturer’s guidelines. RNA samples were sequenced by the University of Delaware Sequencing and Genotyping Center at the Delaware Biotechnology Institute. Libraries were prepared using Perkin Elmer NEXTflex Rapid Directional RNA-seq V1 following the manufacturer’s instructions. Library quality analysis was performed by digital droplet PCR. Sequencing was performed on Illumina NextSeq 550 platform using a NextSeq v2.5 75-cycle high output kit as per the manufacturer’s instructions. Raw sequence data was analyzed by the Center for Bioinformatics and Computational Biology Core Facility at the University of Delaware using the established RNAseq analysis pipeline (adapted from Kalari *et al*.[57]). Quality of sequencing data was assessed using FastQC (ver. 0.10.1; Babraham Bioinformatics). Reads were trimmed for quality (Q<30) and to remove poly-A and Illumina sequencing adapters using Trim Galore! (ver. 0.4.4; Babraham Bioinformatics) and reads less than 35bp after trimming were discarded, resulting in 495.5M quality reads (per sample mean: 41.3M; range: 35.36-45.46M). Trimmed reads were aligned to the M. musculus genome (version mm10) using HiSat2[58] (ver. 2.1.0; mean mapping rate 95.5%), mapping metrics were assessed using RseQC[59] (ver. 2.6.1), and gene/exon feature counts were calculated using HTseq[60] (ver. 0.11.0). Pairwise differential expression analysis was performed to identify gene-level features which are significantly up or down-regulated between treatments using EdgeR[61, 62] (ver. 3.28.1) analyzing genes with a CPM (count per million reads) of at least one in three or more samples. Database for Annotation, Visualization and Integrated Discovery (DAVID)[63, 64] was used to perform GO enrichment analysis and functional classifications.

### Statistical analysis

GraphPad Prism 8 (GraphPad Software Inc) was used to perform statistical analysis. All quantitative data are represented as mean ± standard deviation (SD) or standard error of the mean (SEM). Data shown are from representative results from at least two independent experiments, with the exception of RNAseq data. Tukey’s or Dunnett’s multiple-comparison test or Student’s T-test were used to generate p-values in ANOVA multiple comparisons, unless stated otherwise.

## DATA AVAILABILITY

RNAseq data is submitted to Gene Expression Omnibus (GEO) public repository and data can be accessed through accession number GSE161941.

## ACKNOWLEDGEMENTS

The authors acknowledge J.D. Bhavsar and S.W. Polson for Bioinformatics assistance and B. Kingham for RNA sequencing assistance. The authors also thank W. Chen, A. Kunjapur for Western blotting assistance and access to imager in addition to G. Robbins, J. DeSimone for helpful conversations. BioRender.com was used to create sketches.

## AUTHOR CONTRIBUTIONS

B.M.J., C.A.F. designed research; B.M.J. performed experiments; B.M.J., C.A.F. analyzed data and wrote the paper.

## COMPETING INTERESTS

No competing interests exist.

## FUNDING INFORMATION

Research reported in this publication was supported by the National Institutes of Health and the State of Delaware under Award Number P20GM103446 and P20GM104316, as well as a Research Starter Grant in Pharmaceutics from the PhRMA Foundation, and a University of Delaware Research Foundation Award. The content is solely the responsibility of the authors and does not necessarily represent the official views of the National Institutes of Health.

## Notes

https://www.ncbi.nlm.nih.gov/geo/query/acc.cgi?acc=GSE161941

### Competing Interest Statement

The authors have declared no competing interest.

