## Supplemental Information for "Nanoparticle Internalization Promotes the Survival of Primary Macrophages"

**Table of Contents**


| | |
|---|---|
| PEGDA NP Characterization | S3 |
| Endotoxin Evaluation of NPs | S4 |
| BMM Viability in Serum-Free Cultures | S5 |
| Representative BMM Flow Cytometry Gating | S6 |
| BMM Phenotype Following NP Treatment | S7 |
| DAVID Enrichment Analysis of DEGs | S8 |
| Selected Gene Ontology Processes | S10 |
| Apoptosis Quadrant Analysis Quantification | S11 |
| BMM TUNEL Staining | S12 |
| BMM Lysosomal Tracking | S13 |
| Representative Bronchoalveolar and Peritoneal Lavage Flow Cytometry Gating | S14 |
| Lysosomal Tracking of *In Vivo* Macrophages | S15 |
| TUNEL Staining of *In Vivo* Macrophages | S16 |

### PEGDA NP Characterization

A)

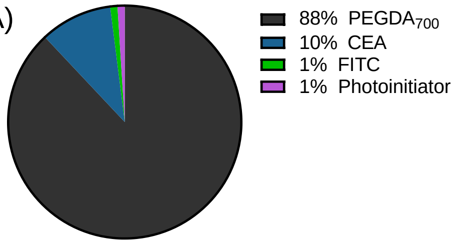


B)

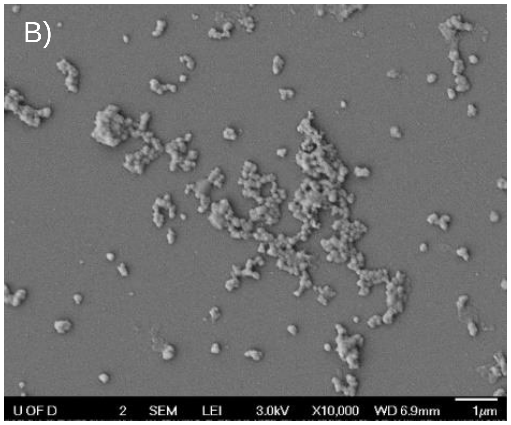


C)

| $D_h$ (nm) | PDI |
|---|---|
| 300 nm | 0.15 |

**Figure S1: PEGDA nanoparticle synthesis and characterization.** A) Synthetic composition of PEGDA nanoparticles B) SEM imaging of PEGDA nanoparticles C) Average hydrodynamic diameter and polydispersity index (PDI) of PEGDA nanoparticles as determined by DLS.

**Endotoxin Evaluation of NPs**

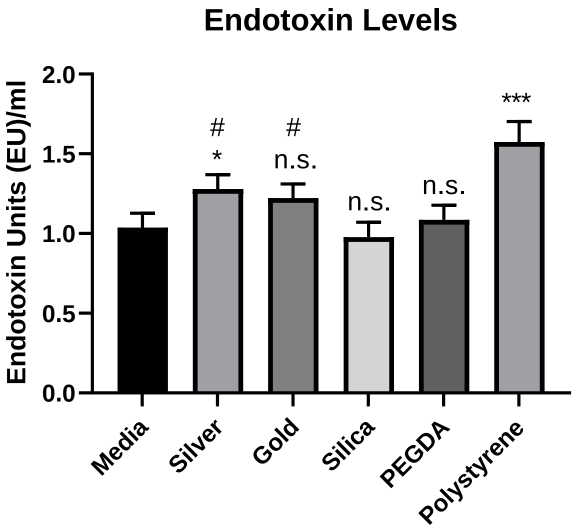


**Figure S2: Endotoxin presence in NP stocks.** Endotoxin units (EU) per ml of NP stocks at a concentration of 100 µg/ml). Statistical analysis performed using Dunnett's multiple comparisons test as part of a one-way ANOVA (*p<0.05, ***p<0.001, n.s. is not significant). Bars represent the mean and error bars represent standard deviation (n=3). Samples labeled with (#) may have significant inherent absorbance that inflates absorbance-detected endotoxin levels.

**BMM Viability in Serum-Free Cultures**

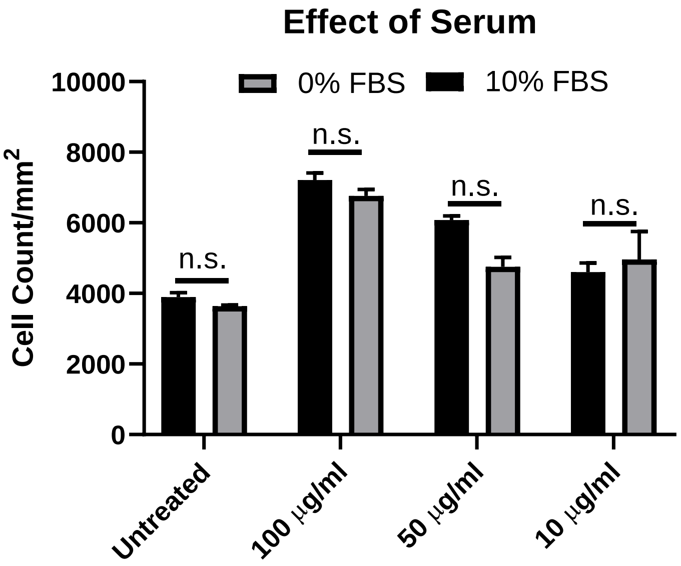


**Figure S3: Effect of fetal bovine serum (FBS) presence on NP-induced BMM survival.** Cell counts of BMMs treated with different concentrations of PEGDA NPs in the presence or absence of serum. Statistical analysis performed using Sidak's multiple comparisons test as part of a two-way ANOVA (n.s. is not significant). Bars represent the mean and error bars represent standard error (n=3).

### Representative BMM Flow Cytometry Gating

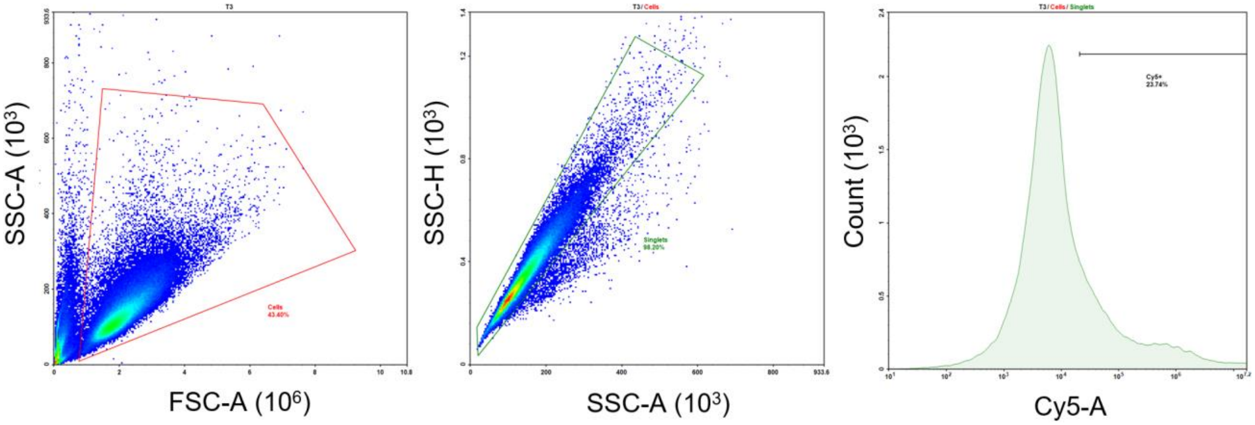


**Figure S4: Representative Flow Cytometry Gating.** Events are gated to remove debris and isolate singlet populations. Cy5+ gating was utilized to determine %NP+ cells.

**BMM Phenotype Following NP Treatment**

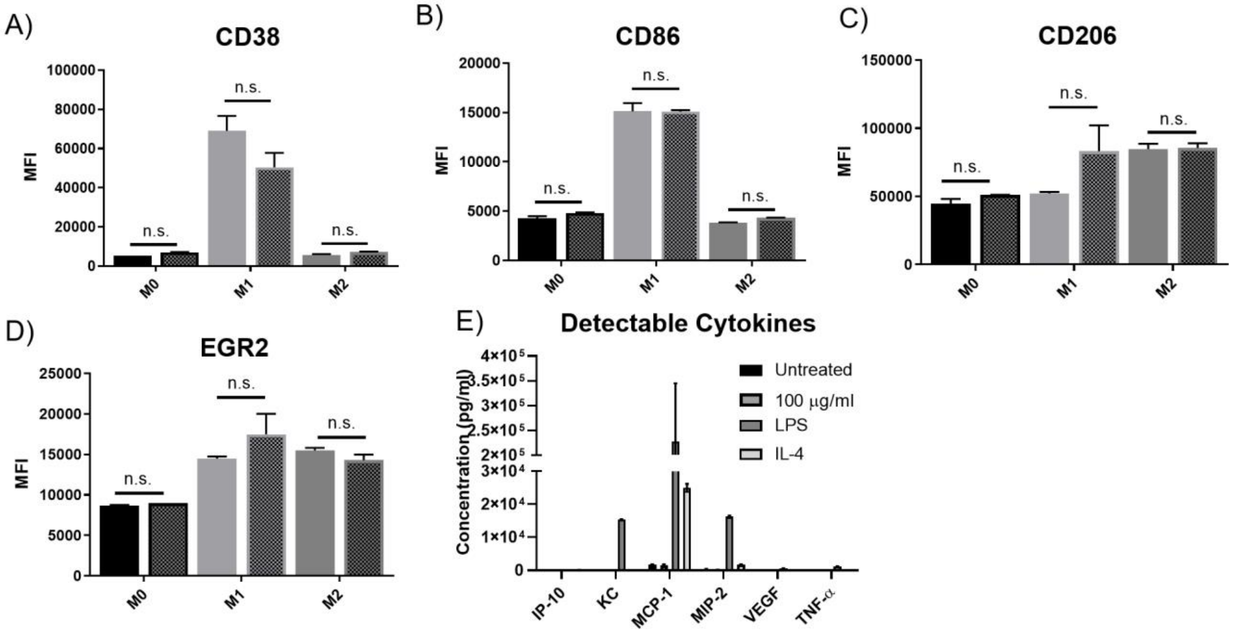


**Figure S5: Phagocytosis of PEGDA NPs retains the initial macrophage phenotype irrespective of polarization and promotes prolonged functionality.** A) CD38 and B) CD86 expression of two representative M1 markers in M0, M1, and M2 BMMs 24 hours following NP treatment with 100 μg/ml NPs (patterned bars). C) CD206 and D) EGR2 expression of two representative M2 marker expression in M0, M1, and M2 BMMs treated as in A. E) Detectable cytokines in supernatants of BMMs with or without NP treatment with LPS and IL-4 as M1 and M2 controls, respectively. In all plots, bars represent the mean and error bars represent SEM (*N*=3). n.s. is not significant using student's T-test. Data in A-D are representative of two independent experiments.

**Table S1: DAVID enrichment analysis of GO Terms in DEGs from RNAseq Analysis.**

| GO Term | GO Type | P-value | Gene Count |
|---|---|---|---|
| *Enrichment Score* 3.25 | | | |
| Positive Regulation of Apoptotic Process | BP | 2.60E-05 | 18 |
| Apoptotic Process | BP | 1.90E-03 | 20 |
| Positive Regulation of Apoptotic Signaling Pathway | BP | 3.30E-03 | 5 |
| *Enrichment Score* 2.07 | | | |
| Endoplasmic Reticulum Membrane | CC | 1.10E-03 | 23 |
| Endoplasmic Reticulum | CC | 4.00E-02 | 29 |
| *Enrichment Score* 2.03 | | | |
| Lysosomal Membrane | CC | 2.60E-06 | 16 |
| Lysosome | CC | 1.50E-03 | 14 |
| *Enrichment Score* 1.92 | | | |
| Immune System Process | BP | 3.30E-03 | 15 |
| *Enrichment Score* 1.63 | | | |
| Cholesterol Homeostasis | BP | 3.20E-03 | 6 |
| Cholesterol Transport | BP | 3.90E-02 | 3 |
| *Enrichment Score* 1.58 | | | |
| Nucleus | CC | 1.90E-04 | 120 |
| Positive Regulation of Transcription, DNA-Templated | BP | 8.90E-04 | 21 |
| Protein Heterodimerization Activity | MF | 1.10E-03 | 19 |
| Negative Regulation of Transcription from RNA Polymerase II Promoter | BP | 1.40E-03 | 24 |
| Transcriptional Repressor Complex | CC | 1.70E-03 | 6 |
| Double-Stranded DNA Binding | MF | 5.60E-03 | 8 |
| Nucleoplasm | CC | 8.90E-03 | 43 |
| Transcription Regulatory Region DNA Binding | MF | 3.10E-02 | 9 |
| Chromatin Binding | MF | 3.20E-02 | 14 |
| DNA Binding | MF | 3.20E-02 | 40 |
| Transcription Corepressor Activity | MF | 3.70E-02 | 7 |
| Positive Regulation of Transcription from RNA Polymerase II Promoter | BP | 4.50E-02 | 24 |
| *Enrichment Score* 1.51 | | | |
| Positive Regulation of Protein Kinase B Signaling | BP | 4.40E-03 | 7 |
| Secretory Granule | CC | 2.70E-02 | 6 |
| *Enrichment Score* 1.48 | | | |
| MAPK Cascade | BP | 7.70E-04 | 7 |
| Negative Regulation by Host of Viral Transcription | BP | 7.70E-04 | 4 |
| Positive Regulation of Phagocytosis | BP | 9.40E-04 | 6 |
| Cellular Response to Organic Cyclic Compound | BP | 1.20E-03 | 7 |
| Protein Kinase B Signaling | BP | 3.00E-03 | 5 |
| Positive Regulation of Protein Kinase B Signaling | BP | 4.40E-03 | 7 |
| Response to Cholesterol | BP | 6.60E-03 | 3 |
| Positive Regulation of ERK1 and ERK2 Cascade | BP | 1.00E-02 | 9 |
| Positive Regulation of Inflammatory Response | BP | 1.80E-02 | 5 |
| Cell Activation | BP | 2.00E-02 | 3 |

| | | | |
|---|---|---|---|
| Positive Regulation of Gene Expression | BP | 2.60E-02 | 13 |
| Immune Response | BP | 3.00E-02 | 10 |
| Inflammatory Response | BP | 4.80E-02 | 11 |
| *Enrichment Score* 1.44 | | | |
| Transport | BP | 5.00E-03 | 44 |
| *Enrichment Score* 1.4 | | | |
| External Side of Plasma Membrane | CC | 8.50E-03 | 12 |
| Cell Surface | CC | 1.40E-02 | 18 |
| *Enrichment Score* 1.39 | | | |
| Positive Regulation of Cell Migration | BP | 1.50E-03 | 1.10E+01 |
| *Enrichment Score* 1.39 | | | |
| Regulation of Cell Proliferation | BP | 2.80E-02 | 9 |
| Immune Response | BP | 3.00E-02 | 10 |
| *Enrichment Score* 1.32 | | | |
| Positive Regulation of Cell Proliferation | BP | 1.60E-04 | 22 |
| Negative Regulation of Apoptotic Process | BP | 1.90E-02 | 17 |
| Positive Regulation of Pri-miRNA Transcription From RNA Polymerase II Promoter | BP | 4.70E-02 | 3 |
| *Enrichment Score* 1.26 | | | |
| Transforming Growth Factor Beta Receptor Signaling Pathway | BP | 6.70E-03 | 6 |
| JNK Cascade | BP | 1.60E-02 | 4 |
| Positive Regulation of JUN Kinase Activity | BP | 2.50E-02 | 4 |
| Activation of MAPK Activity | BP | 3.20E-02 | 5 |
| *Enrichment Score* 1.2 | | | |
| Response to Drug | BP | 8.00E-03 | 13 |
| Aging | BP | 2.00E-02 | 8 |
| *Enrichment Score* 1.15 | | | |
| Cellular Response to Tumor Necrosis Factor | BP | 3.10E-02 | 6 |
| *Enrichment Score* 1 | | | |
| Histone Binding | MF | 4.50E-02 | 6 |
| *Enrichment Score* 1 | | | |
| Metalloaminopeptidase Activity | MF | 3.50E-02 | 3 |
| Peptide Catabolic Process | BP | 4.70E-02 | 3 |
| *Enrichment Score* 0.96 | | | |
| Cell Migration | BP | 9.50E-04 | 11 |
| *Enrichment Score* 0.92 | | | |
| Ubiquitin-Protein Transferase Activity | MF | 3.30E-02 | 11 |
| *Enrichment Score* 0.84 | | | |
| Kinase Activity | MF | 2.00E-02 | 19 |
| Nucleotide Binding | MF | 4.10E-02 | 41 |
| Protein Phosphorylation | BP | 4.10E-02 | 16 |
| *Enrichment Score* 0.81 | | | |
| Membrane | CC | 2.40E-07 | 148 |
| *Enrichment Score* 0.37 | | | |
| Poly(A) RNA Binding | MF | 4.40E-02 | 26 |

### Selected Gene Ontology Processes

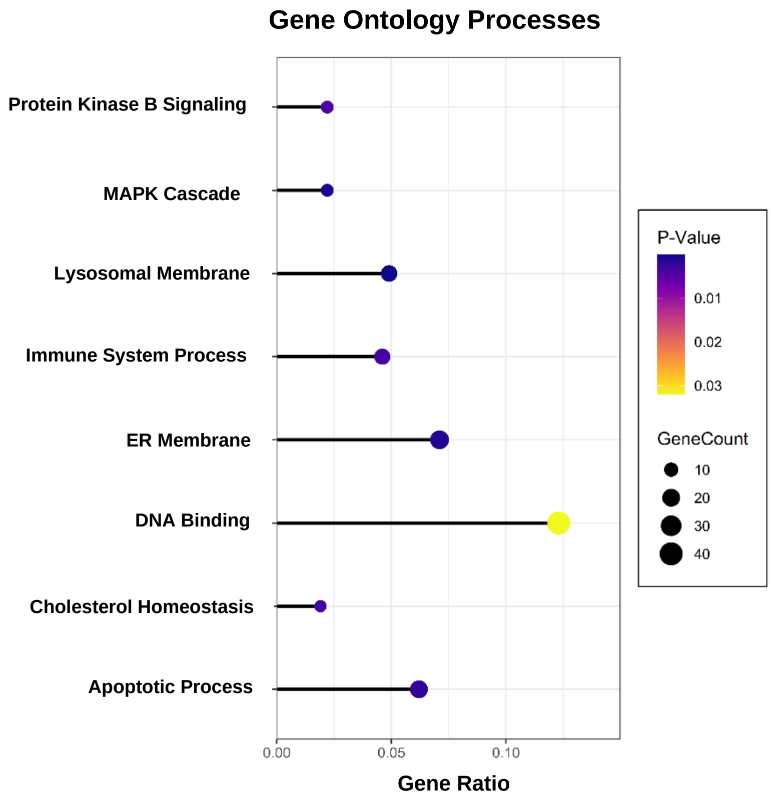


**Figure S6: Selected statistically significant Gene Ontology (GO) terms** from functional annotation clustering of top 331 differentially expressed genes (DEGs) using DAVID Tool

**BMM Apoptosis Quadrant Analysis Quantification**

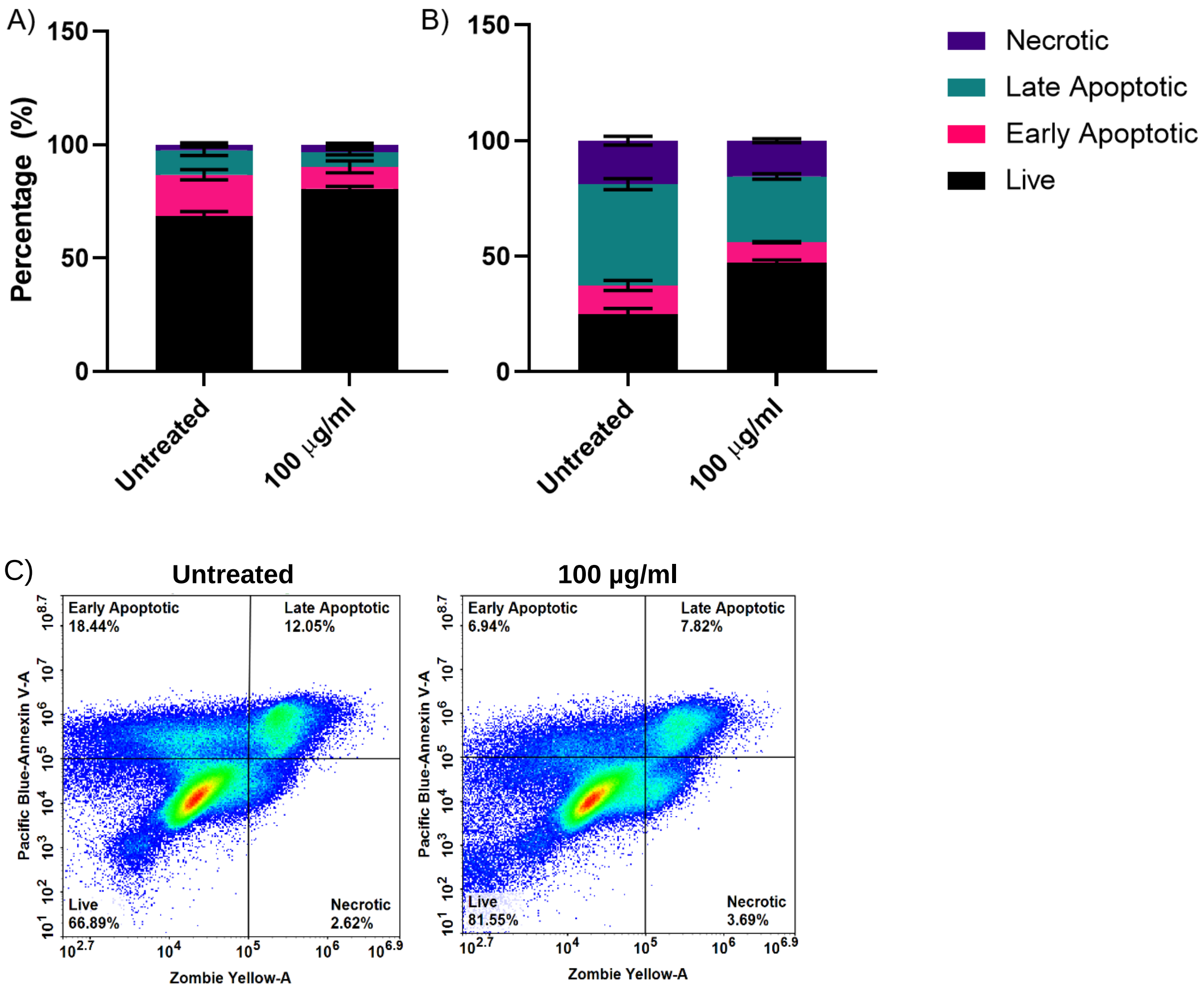


**Figure S7: Quantitative results of apoptosis quadrant analysis.** Percentages of Live, Early Apoptotic, Late Apoptotic, and Necrotic populations of BMMs A) 24 hours B) 72 hours following NP treatment. C) Representative flow quadrant gating.

### BMM TUNEL Staining

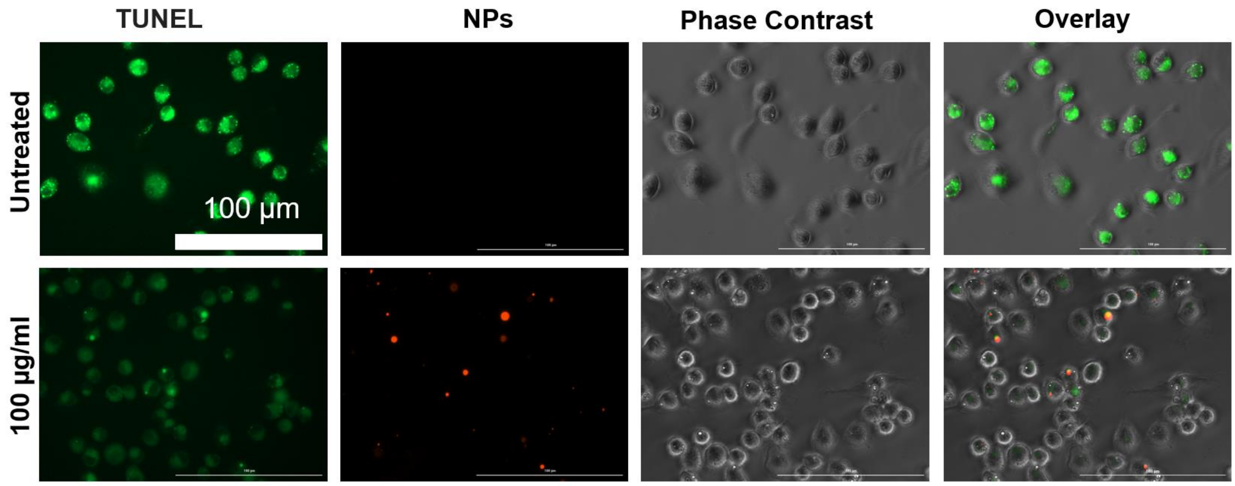


**Figure S8: TUNEL apoptosis imaging analysis of untreated or NP-treated BMMs 72 hours following NP treatment at 40x magnification.**

**BMM Lysosomal Tracking**

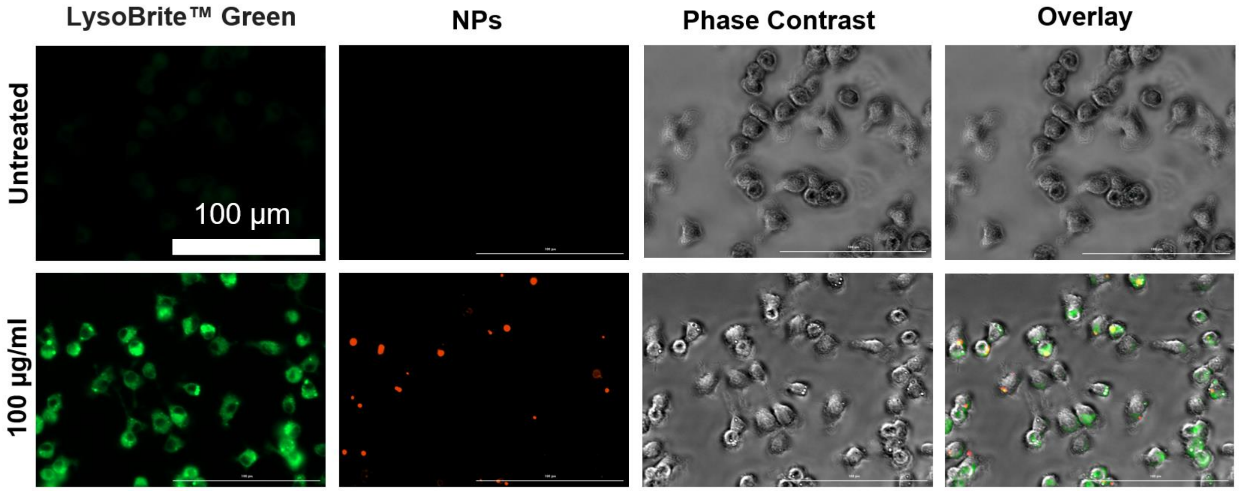


**Figure S9: Lysosomal tracking and imaging at 40x magnification with LysoBrite™ Green 72 hours following NP treatment.**

### Representative Bronchoalveolar and Peritoneal Lavage Flow Cytometry Gating

A)

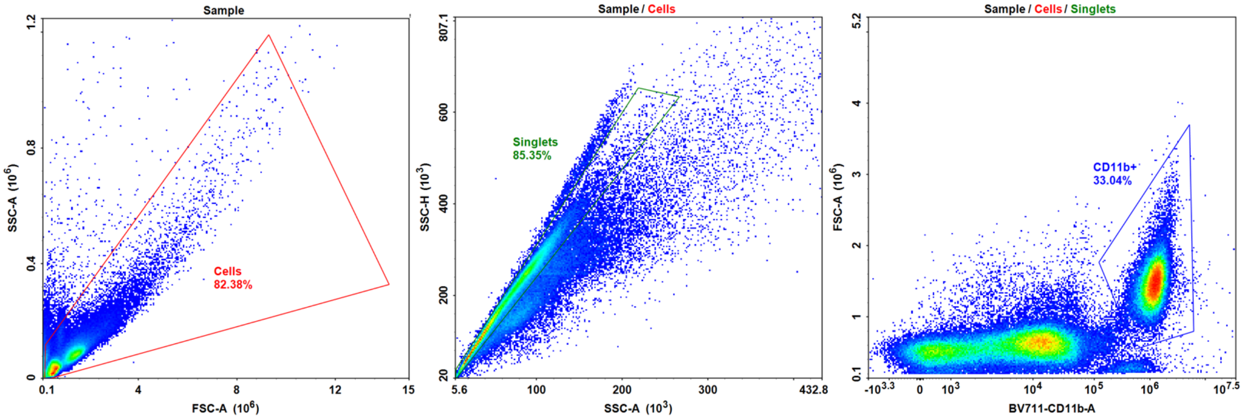


B)

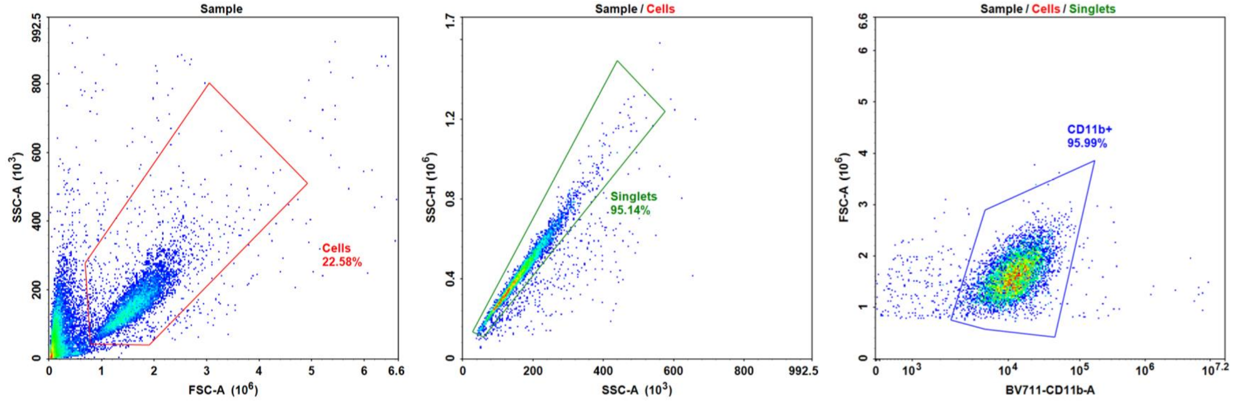


**Figure S10: Representative Flow Cytometry Gating of A) Peritoneal Lavage and B) Bronchoalveolar Lavage.** Cells in the lavages were separated from debris and single cell populations were identified. CD11b+ macrophages were gated for subsequent analyses.

**Lysosomal Tracking of *In Vivo* Macrophages**

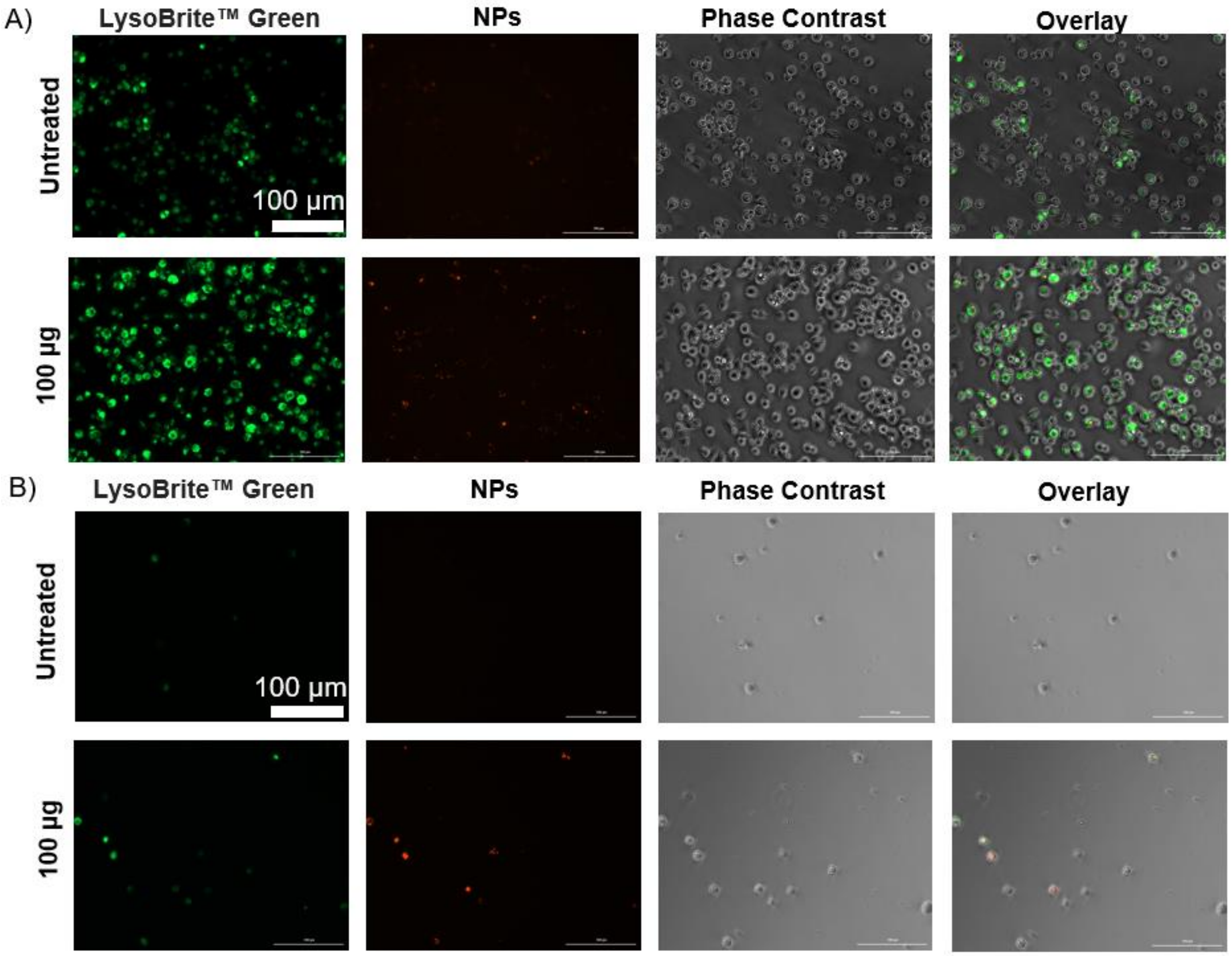


**Figure S11: Lysosomal Tracking.** Fluorescent imaging at 20x magnification with LysoBrite™ Green in A) Peritoneal and B) Alveolar macrophages 24 hours following *in vivo* dosing.

**TUNEL Staining of *In Vivo* Peritoneal Macrophages**

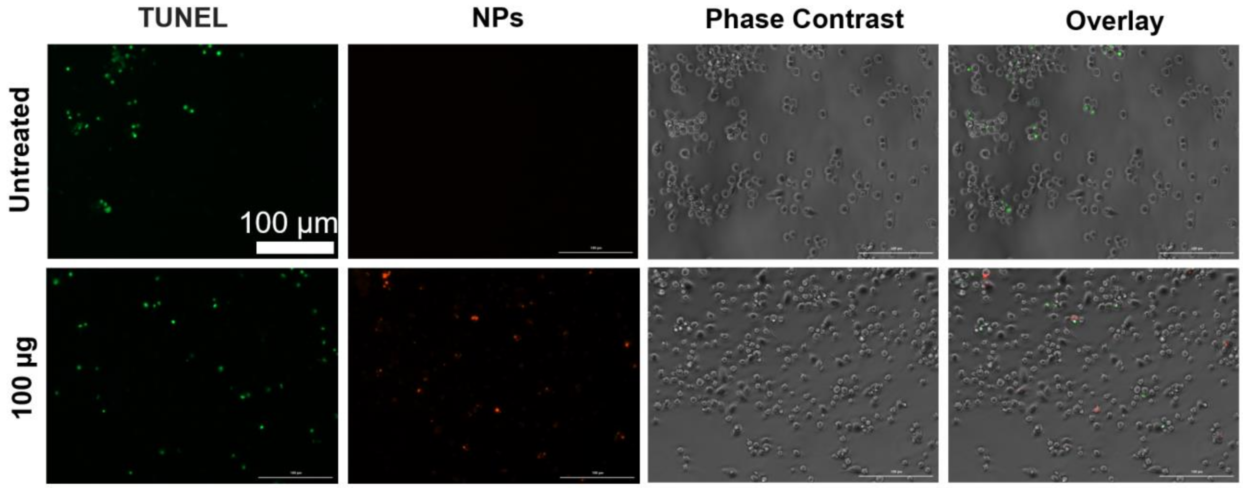


**Figure S12: TUNEL Apoptosis Staining.** Fluorescent imaging at 20x magnification with TUNEL in Peritoneal macrophages 24 hours following *in vivo* dosing.
